# A Novel and Prevalent Pathway in Microbial Selenium Metabolism

**DOI:** 10.1101/2022.04.13.486033

**Authors:** Chase M. Kayrouz, Mohammad R. Seyedsayamdost

**Affiliations:** Department of Chemistry, Princeton University, Princeton, NJ 08544, United States; Department of Molecular Biology, Princeton University, Princeton, NJ 08544, United States

## Abstract

Selenium is an essential micronutrient in diverse organisms. Two routes are known for its insertion into proteins and nucleic acids via selenocysteine and 2-selenouridine, respectively^1^. However, despite its importance, pathways for specific incorporation of selenium into small molecules have remained elusive. We herein use a genome mining strategy to uncover a widespread three-gene cluster in varied microorganisms that encodes a dedicated pathway for producing selenoneine, the selenium-analog of the multifunctional molecule ergothioneine^2,3^. We elucidate the reactions of all three proteins and uncover two novel selenium-carbon bond-forming enzymes and the first biosynthetic pathway for production of a selenosugar, an unexpected intermediate on route to the final product. Our findings expand the scope of biological selenium utilization, suggest that the selenometabolome is more diverse than previously thought, and set the stage for the discovery of other selenium-containing natural products.

## Main Text

Selenium (Se) is an essential trace element across all kingdoms of life and is found mainly in selenoprotein and selenonucleic acid biopolymers^1^. Selenoproteins carry Se in the form of selenocysteine (Sec), which often constitutes the redox active center in the catalytic reduction of peroxides, disulfides, sulfoxides, and aryl iodides^4–8^. Selenonucleic acids carry Se in 5-methylaminomethyl-2-selenouridine, which is incorporated into the wobble position of several tRNAs by a variety of microbes^9^. Aside from its presence in important biopolymers, Se has also been detected in place of sulfur in many common sulfur-containing metabolites, such as methionine, cystathionine, and dimethyl sulfide, when organisms are grown under Se-enriched conditions. However, sulfur and Se are poorly discriminated by biosynthetic enzymes, and thus the observed selenometabolites are usually nonspecific products of sulfur-utilizing enzymes^10^. Se-specific biosynthetic pathways have so far been limited exclusively to Sec and selenouridine; they encompass a grand total of three Se-specific enzymes. Common to both pathways is selenophosphate synthetase (SelD), which phosphorylates hydrogen selenide to generate selenophosphate (SeP), the substrate for selenocysteine synthase (SelA) and selenouridine synthase (SelU, **Fig. 1a**). Using SeP, SelA converts seryl-tRNA_Sec_ into selenocysteinyl-tRNA_Sec_ in a pyridoxal-5-phosphate (PLP)-dependent fashion, while the rhodanese enzyme SelU catalyzes a S-to-Se substitution on 2-thiouridine-containing tRNAs^11–13^ (**Fig. 1a**).

**Figure 1.**
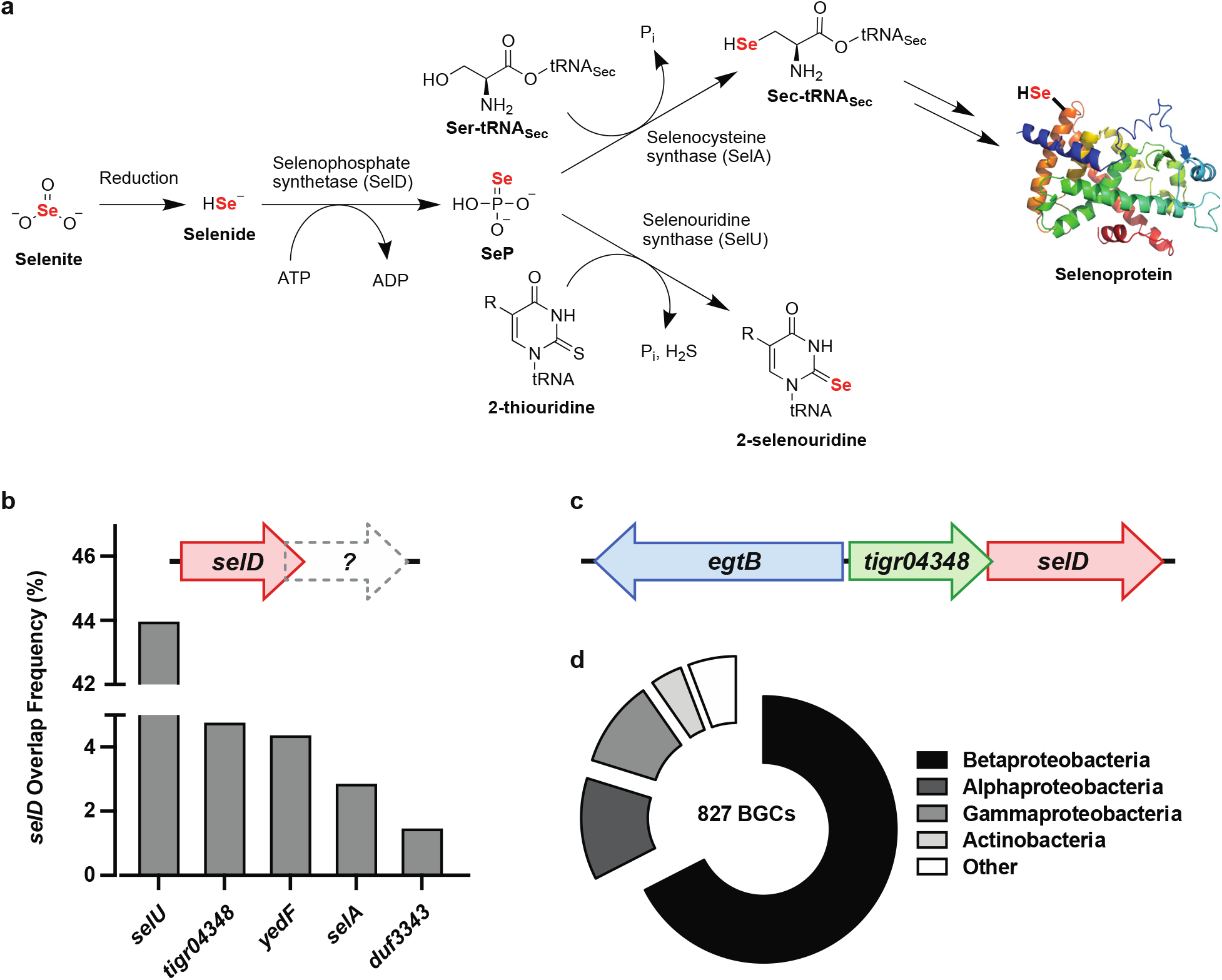
Known and new biological pathways for Se incorporation. **a**, Pathway for insertion of Se into protein (top) and nucleic acid biopolymers (bottom). **b**, Distribution of genes that overlap with all known *selD* sequences in the available genome database. TIGR04348, a previously uncharacterized protein family, is the second most common. **c**, A new potential Se-insertion BGC consisting of an *egtB* homolog, *tigr04348*, and *selD*. **d**, Prevalence of the three-gene cassette in panel **c**.

Given the taxonomic ubiquity of Se utilization, it is hard to imagine that Se would be limited to these two pathways. Furthermore, both SelA and SelU, the only known C-Se bond-forming enzymes, act on macromolecular substrates, thus leaving no known mechanism for specific incorporation of Se into organic small molecules, notably natural products, which can harbor unusual atoms and complex architectures. We therefore speculated that Nature may have indeed evolved additional biosynthetic strategies for endowing small molecules with Se and its unique properties. Guided by a bioinformatic search, we herein report a novel and widespread pathway for incorporation of Se into microbial metabolites, involving two novel and unusual Se-C bond-forming enzymes.

### Genome mining for selenometabolite pathways

To identify new selenometabolite pathways, we first envisioned potential genetic characteristics of organisms that may encode them. Given that SeP is the common Se-donor to both selenoproteins and selenonucleic acids, we hypothesized that new pathways may follow a similar biosynthetic trajectory. Conveniently, genes involved in the production of natural products are often grouped into biosynthetic gene clusters (BGCs) in microbial genomes, thereby allowing us to formulate a basic criterion for genome mining: the presence of *selD* within a BGC^14^.

We retrieved all *selD* genes from the National Center for Biotechnology Information (NCBI) database and, after a strict dereplication process, extracted the genetic contexts of each *selD* gene to search for the presence of common natural product biosynthetic genes. Unfortunately, we were unable to identify obvious instances of *selD* within a natural product BGC. We therefore inverted the search to interrogate *selD* genetic contexts in a hypothesis-free manner and identify genes commonly co-localized with *selD*. Specifically, we quantified the abundance of genes, which overlap with a *selD* open-reading-frame by one or more base-pairs, reasoning that this additional constraint would select for genes whose products are transcriptionally linked to SelD (**Fig. 1b**). Among the top five *selD*-overlapping genes are *selA* and *selU*, validating the strategy’s ability to identify functionally-related genes, along with two genes, *yedF* and *duf3343*, that are thought to serve an as-yet-unidentified role in Se reduction and/or trafficking^15^.

The second most commonly-overlapping gene, annotated as a putative glycosyltransferase from the TIGR04348-family, caught our attention (**Fig. 1b, Supplementary Table 1**). Though accounting for nearly 5% of all *selD*-overlapping genes, to the best of our knowledge, TIGR04348 has never been implicated in Se metabolism, nor has any member of its protein family been characterized. Furthermore, a closer examination of *selD*-*tigr04348* genetic contexts revealed a third commonly co-localized gene: a homolog of *egtB*, which encodes the C-S bond-forming enzyme in ergothioneine biosynthesis^16^ (**Fig. 1c**). Intriguingly, over 800 examples of this three-gene cassette were found among diverse bacterial genomes, suggesting that it may encode a widespread pathway dedicated to the biosynthesis of a new selenometabolite (**Fig. 1d**). This hypothesis is further endorsed by the presence of an additional gene coding for a putative selenate transporter in close proximity to many of the retrieved clusters (**Supplementary Fig. 1**). We therefore set out to characterize this pathway using metabolomic and biochemical approaches.

### Microbial metabolite profiling

We began by performing a metabolomic analysis of two species that harbor the *selD-egtB-tigr04348* cluster. The actinomycete *Amycolatopsis palatopharyngis* DSM 444832 and the β-proteobacterium *Variovorax paradoxus* DSM 30034 were grown in the presence of sodium selenite (**Supplementary Fig. 1**). The cells were then pelleted and analyzed by HPLC-MS, revealing the production of ergothioneine and its Se-analog, selenoneine, by both bacteria (**Fig. 2a, 2b, Supplementary Table 2**). Interestingly, selenoneine has been observed on several occasions in biological samples, with particularly high abundance in the blood and tissues of marine animals^2,17^. The natural origin of selenoneine has thus far remained enigmatic, though researchers have speculated that it may arise from non-specific incorporation by the ergothioneine pathway^18^. Indeed, upon further genomic examination, both strains were found to harbor a canonical ergothioneine BGC (*egt*) in addition to the putative selenometabolite BGC. To examine whether selenoneine may be the product of nonspecific Se incorporation by the *egt* cluster, two close relatives of the producing strains, the actinomycete *Streptomyces rimosus* ATCC 10970 and the β-proteobacterium *Burkholderia thailandensis* E264, which encode a canonical *egt* cluster but lack the putative selenometabolite cassette, were analyzed in the same fashion, revealing exclusive production of ergothioneine, and not selenoneine (**Fig. 2a**). These observations are consistent with prior reports of the inability of the bacterial ergothioneine biosynthetic machinery to generate selenoneine in vitro^19^. Our results suggest that selenoneine may in fact be the product of the new cluster, which we have termed *sen*, with *senA, senB*, and *senC* coding for an *egtB* homolog, a putative glycosyltransferase, and a *selD* homolog, respectively (**Fig. 2c**).

**Figure 2.**
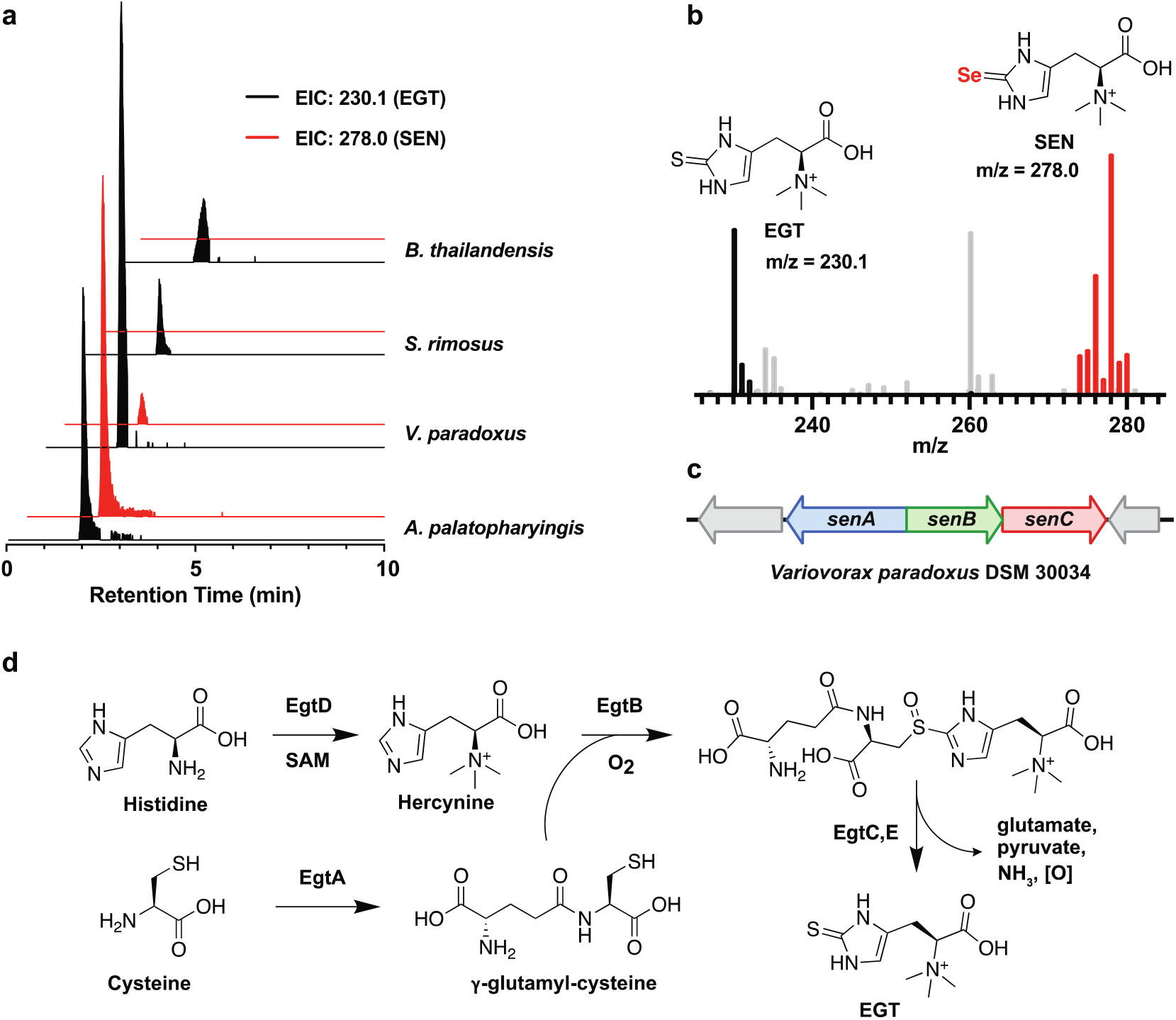
Production of selenoneine by bacteria that carry the three-gene *sen* cluster. **a**, Production of selenoneine (SEN, *m/z* 278.0) and ergothioneine (EGT, *m/z* 230.1), determined by HPLC-MS analysis, by the strains indicated. Only those that have the *sen* cluster synthesize selenoneine. **b**, HR-MS spectra of *A. palatopharyngis* culture extracts, revealing production of both ergothioneine and selenoneine. **c**, The three-gene *sen* cluster consisting of the *egtB* homolog *senA*, a member of an uncharacterized superfamily *senB*, and a *selD* homolog *senC*. **d**, Ergothioneine biosynthetic pathway.

If correct, our hypothesis would indicate two distinct biosynthetic routes to ergothioneine and selenoneine. Ergothioneine is produced by *egtABCDE*. Early steps involve EgtA-catalyzed isopeptide bond formation between cysteine and glutamic acid, and S-adenosylmethionine (SAM)-dependent trimethylation of histidine by EgtD (**Fig. 2d**). The products, γ-glutamyl-cysteine (GGC) and hercynine, are then oxidatively linked via a sulfoxide by the atypical nonheme iron enzyme EgtB. Finally, following EgtC-catalyzed hydrolysis of the glutamic acid residue, EgtE performs PLP-dependent C-S bond cleavage to furnish ergothioneine^16^. In contrast, the *sen* cluster contains only three genes. We speculated that SenB utilizes SeP (generated by SenC) to generate a new selenometabolite. SenA could then catalyze C-Se bond-formation between this new species and hercynine, followed by C-Se bond-cleavage to give selenoneine. Hercynine would likely be siphoned from the canonical ergothioneine pathway, though a small subset of *sen* clusters feature a co-localized *egtD* gene (**Supplementary Fig. 1**).

### Reconstitution of selenoneine biosynthesis

To evaluate this proposal and conclusively link the production of selenoneine to the *sen* cluster, we reconstituted the entire biosynthetic pathway in vitro. SenA, SenB, SenC, and EgtD were cloned from *V. paradoxus* and isolated as soluble, His-tagged proteins through heterologous expression in *E. coli* (**Supplementary Tables 3-4**). As expected, SenC was found to catalyze the ATP-dependent phosphorylation of sodium selenide to yield SeP (**Fig. 3a, Supplementary Fig. 2**). Next, we evaluated the function of SenB, the C-terminus of which shows weak homology to NDP-sugar binding domains of glycosyltransferases. When SenB was added to the SenC reaction along with the common sugar donor UDP-glucose, production of a new species was observed. It contained two Se atoms, as judged by the characteristic Se isotope pattern obtained by HR-MS, implying a potential diselenide-dimer (**Fig. 3b**). Derivatization of the species with the thiol-labeling reagent monobromobimane (mBBr) resulted in an mBBr-derivative with a single Se atom and a mass consistent with the monomeric form of the underivatized species (**Fig. 3c**). Purification of the mBBr-derivative and structural elucidation by multidimensional NMR spectroscopy revealed 1-seleno-β-D-glucose (SeGlc), a rare example of a naturally-occurring selenosugar, as the SenB reaction product (**Fig. 3d, Supplementary Fig. 3, Supplementary Table 5**).

**Figure 3.**
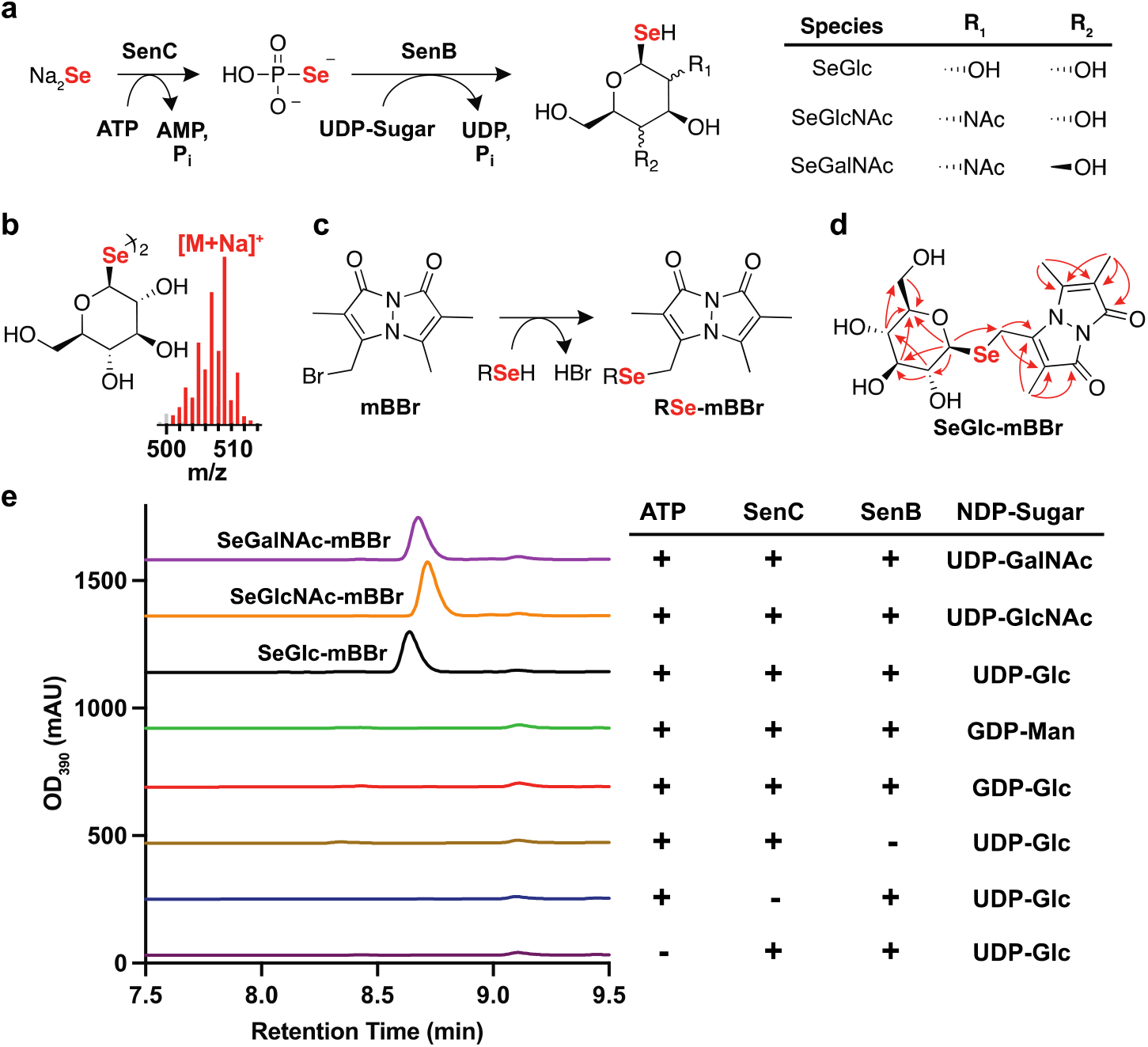
SenB, a novel selenosugar synthase. **a**, Reactions shown to be carried out by SenC (SelD homolog) and SenB, the first selenosugar synthase. SenB accepts three different UDP-sugars indicated to generate selenoglucose (SeGlc), seleno-N-acetylglucosamine (SeGlcNAc), or seleno-N-acetylgalactosamine (SeGalNAc). **b**, Diselenide product of the SenB reaction. **c**, The mBBr derivatization reaction. **d**, Structure of the mBBr-derivatized SeGlc; relevant ^1^H-^13^C HMBC NMR correlations (red arrows) used to solve the structure are shown. **e**, Substrate tolerance and control reactions for SenB. The reaction content for each trace is shown in the table to the right of the HPLC-UV chromatogram. The product peaks in the chromatogram are labeled.

Control assays further confirmed the requirement of each reaction component (**Fig. 3e**). In the absence of SenB, no SeGlc was formed. Controls lacking either SenC or ATP also abolished SeGlc production, verifying SeP as the source of Se for SenB. Moreover, no product was observed with sodium sulfide, suggesting SenC discriminates between Se and S, thus synthesizing only SeP for the downstream SenB-catalyzed reaction (**Supplementary Fig. 2**). When assayed against various nucleotide-sugar substrates, SenB was found to efficiently utilize UDP-N-acetylglucosamine (UDP-GlcNAc) and UDP-N-acetylgalactosamine (UDP-GalNAc, **Supplementary Fig. 4-5, Supplementary Table 6**), but not GDP-mannose or GDP-glucose, suggesting SenB is specific for UDP-sugars (**Fig. 3e**). Together, these results allow us to designate SenB as a novel selenosugar synthase, which now joins SelA and SelU as only the third bona fide Se-C bond-forming enzyme characterized to date.

**Figure 4.**
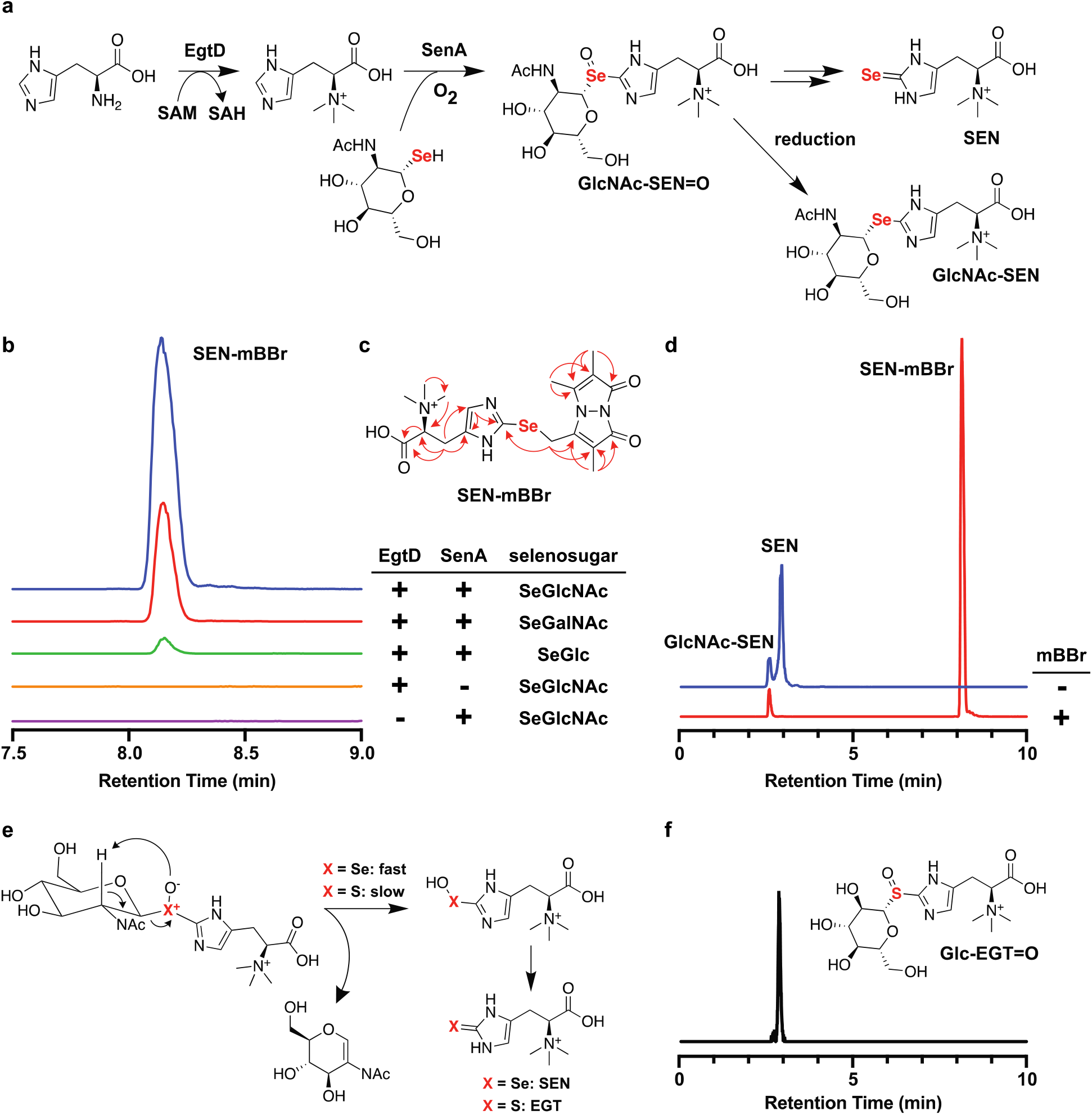
SenA, a novel selenoneine synthase. **a**, Biosynthetic pathway for selenoneine (SEN) with the reactions elucidated in this work. Hercyncyl-SeGlcNAc selenoxide (GlcNAc-SEN=O) can spontaneously convert to selenoneine (see panel **e**) or be reduced to hercynyl-SeGlcNAc seleno-ether (GlcNAc-SEN). **b**, Reaction of SenA monitored by HPLC-MS. The reaction content for each trace in the chromatogram is shown in the table to the right. Among sugars tested, SenA shows preference for SeGlcNAc. **c**, Structure of SEN-mBBr. Relevant ^1^H-^13^C HMBC correlations are shown (red arrows). **d**, Selenoneine can be detected directly by HR-MS in the absence of mBBr derivatization. **e**, Proposed internal Cope elimination to form selenoneine. Cope eliminations are significantly slower when Se is replaced with S. **f**, HPLC-MS detection of hercynyl-thioglucose sulfoxide (Glc-EGT=O) when the SenA reaction is carried out with a thiosugar donor.

The remaining enzyme encoded in the *sen* cluster, SenA, is a distant homolog of the C-S bond-forming sulfoxide synthase EgtB^16^. Previous work by the groups of Seebeck and Liu has provided extensive characterization of this family of nonheme iron enzymes, including crystal structures that have pinpointed residues involved in iron-, hercynine-, and thiol-binding^20–22^. SenA proteins bear 30-50% similarity to members of the EgtB family and share its conserved iron-binding three-His facial triad, catalytic Tyr residue, and motifs involved in hercynine-binding (**Supplementary Fig. 6**). However, a conserved Arg87/Asp416 pair implicated in thiol binding within the EgtB active site is replaced with His and Phe or Tyr, respectively, in all SenA proteins. These substitutions suggested to us that SenA catalyzes Se-C bond-formation between hercynine and a different substrate, presumably a selenosugar, en route to selenoneine. We tested this hypothesis by first recapitulating the activity of *V. paradoxus* EgtD in vitro, allowing for rapid formation of hercynine from histidine and SAM (**Fig. 4a, Supplementary Fig. 7**). Upon incubation of SenA with hercynine (generated in situ with EgtD, His, and SAM), various selenosugars, dithiothreitol (DTT), and Fe(II), followed by derivatization with mBBr, we observed a new species containing a single Se atom and a high-resolution mass consistent with that of selenoneine-mBBr (**Fig. 4b**). The structure was confirmed by NMR spectroscopy upon purification of the product from large-scale reactions with SeGlcNAc (**Fig. 4c, Supplementary Fig. 8, Supplementary Table 7**), the substrate for which SenA showed the greatest preference. More importantly, in the absence of mBBr, selenoneine itself was directly observed in the product mixture as evidenced by HR-MS and HR-MS/MS analysis (**Supplementary Table 8, Supplementary Fig. 9**), thus completing reconstitution of the entire selenoneine biosynthetic pathway in vitro and confirming that the *sen* cluster is responsible for building this unusual natural product (**Fig. 4d**).

In ergothioneine biosynthesis, the EgtB reaction yields hercynyl-GGC sulfoxide, which is further processed by EgtC and EgtE to produce ergothioneine^23,24^ (**Fig. 2d**). The analogous product by SenA, hercynyl-SeGlcNAc selenoxide, was not detected in our reaction. However, further examination of the mBBr-derivatized reaction revealed the reduced form of this product, hercynyl-SeGlcNAc selenoether; its structure was corroborated by HR-MS and HR-MS/MS analysis (**Fig. 4d, Supplementary Table 8, Supplementary Fig. 9**). We suspect that this product forms as a result of reduction of the selenoxide intermediate by excess DTT in the reaction, implying that the selenoxide intermediate is indeed formed. Spontaneous selenoxide eliminations are well-documented, and this intermediate could undergo, aside from reduction by DTT, rapid internal Cope elimination to form the selenenic acid (**Fig. 4e**), which would convert to selenoneine through either reduction or disproportionation^25^. Selenoxide internal eliminations occur upwards of 10^5^ times more rapidly than the related sulfoxide eliminations^1^. Thus, the tendency of selenoxides to undergo internal elimination obviates the need for an EgtE-like reaction in selenoneine biosynthesis. In support of this proposal, replacement of SeGlcNAc with commercially available thioglucose in the SenA reaction mixture resulted in accumulation of hercynyl-thioglucose sulfoxide, with no observed formation of ergothioneine or reduction to the corresponding thioether (**Fig. 4f, Supplementary Table 8, Supplementary Fig. 9**).

## Conclusions

Our results open new avenues in selenobiology, notably the incorporation of Se into natural products. These molecules are typically comprised of a limited set of elements, usually consisting of C, N, O, H, and S. Demonstration of a widespread pathway for Se incorporation into natural products provides impetus for interrogating other selenometabolites and their biosynthetic origins that are like encoded in microbial genomes. The reactions of SenA (selenoneine synthase) and SenB (selenosugar synthase) reported herein double the number of previously known Se-C-bond forming enzymes in Nature and at the same time stimulate future studies into their reaction mechanisms. Seminal studies by Stadtman and Böck previously demonstrated pathways for incorporation of Se into protein and nucleic acid biopolymers^9,11–13^. SenABC provide the first pathway for biosynthesis of selenosugars, which could play yet unknown roles in primary and secondary metabolism. More generally, selenometabolomes provide an untapped chemical space in bacteria that can now be mined for novel natural products and their associated bioactivities.

## Methods

### Materials

All materials were purchased from Millipore-Sigma or Fisher Scientific unless otherwise specified. Cloning reagents were purchased from New England BioLabs (NEB). Codon-optimized gene fragments were purchased from Genewiz. DNA primers were purchased from Sigma.

### General experimental procedures

High-performance liquid chromatography-coupled mass spectrometry (HPLC-MS) was performed on an Agilent instrument equipped with a 1260 Infinity Series HPLC, an automated liquid sampler, a photodiode array detector, a JetStream ESI source, and the 6540 Series Q-tof mass spectrometer. HPLC purifications were carried on an Agilent 1260 Infinity Series HPLC system equipped with a photodiode array detector and an automated fraction collector. Solvents for all LC-MS and HPLC experiments were water + 0.1% formic acid (Solvent A) and MeCN + 0.1% formic acid (Solvent B). Nuclear magnetic resonance (NMR) spectra were collected at the Princeton University Department of Chemistry’s NMR facility on a Bruker Avance III 500 MHz NMR spectrometer equipped with a DCH double resonance cryoprobe. NMR data were processed and analyzed using MestReNova software. Anaerobic enzyme assays were carried out in an Mbraun glovebox under a N_2_ atmosphere maintained at <0.1 ppm O_2_.

### Bioinformatics

All SelD protein sequences were retrieved from the NCBI database using a combination of bioinformatic tools. First, the NCBI E-Direct toolkit was used to extract all proteins from the conserved domain database (CDD) with a match to the SelD position-specific scoring matrix (PSSM) “COG0709”. This resulted in 205,276 retrieved sequences. Next, the CD-hit clustering tool was used to cluster the sequences at 98% sequence identity to dereplicate and reduce redundancy in the dataset resulting from over-sequencing bias^26^. This reduced the dataset to 31,280 sequences, illustrating the amount of sequence redundancy. These were then aligned against a short list of representative bacterial SelD sequences using the blastp algorithm with an E-value cutoff of 1e-20 to remove any non-SelD protein families, such as thiamine-phosphate kinase, which has slight homology to SelD^27,28^. This resulted in a final, strictly-dereplicated set of 10,947 SelD protein sequences from the NCBI database.

Next, the E-Direct toolkit was used to extract all annotated open-reading-frames (ORFs) within a window 4-kbp upstream or downstream of each *selD* gene. From this dataset, a total of 2960 ORFs with a minimum of 1-bp overlap with a *selD* ORF were obtained and re-annotated using a combination of the NCBI CDD-search tool and manual examination. The protein families corresponding to the ten most common *selD*-overlapping genes are shown in Supplementary Table 1.

To further examine the *tigr04348* gene, first a blastp search was performed against the NCBI non-redundant protein database using a short list of TIGR04348 protein sequences from various phyla and an E-value cutoff of 1e-20. This resulted in a list of 2,353 TIGR04348 sequences after dereplication by strain name. Next, the E-Direct toolkit was used to extract 5-kbp upstream and downstream of each *tigr04348* gene. These genetic regions were then used to construct a BLAST database and interrogated using the tblastn algorithm for the presence of *egtB* and *selD* homologs, resulting in a total of 827 *egtB-tigr04348-selD* BGCs, henceforth referred to as the *sen* BGC consisting of sen*A, senB*, and *senC*. A select few examples of *sen* BGCs are depicted in Supplementary Fig. 1, noting particularly the presence of co-localized *egtD* and a putative selenate transporter gene in a subset of them.

For amino acid multiple sequence alignments of EgtB and SenA, the top 500 SenA sequences most similar to SenA from *V. paradoxus* and the top 4000 EgtB sequences most similar to EgtB from *Mycobacterium thermoresistibile* were retrieved using BLAST and aligned separately using MAFFT with the FFT-NS-2 algorithm^29^. These alignments were then used to generate logo plots of the important motifs, as determined by Seebeck, et al. for *M. thermoresistibile*, using the Logomaker python script^20,30^ (Supplementary Fig. 6).

### Strains, media, and general culture conditions

Strains used in selenometabolite characterization and recombinant protein production experiments are listed in Supplementary Table 2. ISP Medium 2 agar plates and tryptic soy broth were used for general maintenance and liquid cultures of *Amycolatopsis palatopharyngis* DSM 44832 and *Streptomyces rimosus* ATCC 10970. Nutrient agar plates and nutrient broth were used for general maintenance and liquid cultures of *Variovorax paradoxus* DSM 30034. LB agar and broth (supplemented with 50 mM MOPS, pH 7.0) were used for general maintenance and liquid cultures of *Burkholderia thailandensis* E264.

### Selenometabolite production screens

For each strain, a single colony from an agar plate was inoculated into a sterile culture tube containing 5 mL of liquid medium and incubated at 30°C/250 rpm. The starter cultures were then used to inoculate 5 mL liquid cultures supplemented with 100 μM of filter-sterilized Na_2_SeO_3_ and incubated at 30°C/250 rpm. Production cultures of *B. thailandensis* were grown for 24 hours, *V. paradoxus* and *S. rimosus* for 48 hours, and *A. palatopharyngis* for 72 hours. Following incubation, 2 mL of each culture was transferred to an Eppendorf tube, pelleted by centrifugation, and supernatants were removed. Cell pellets were resuspended in 0.4 mL of ddH_2_O and subjected to hot-water extraction by heating to 95°C for 15 minutes. After cooling to room temperature, the mixtures were pelleted by centrifugation and supernatants were analyzed by HPLC-MS. Analytes were separated on a Synergi Fusion-RP column (Phenomenex, 100 × 4.6 mm, 4 µm) with a flow rate of 0.5 mL/min and an elution program consisting of 0% solvent B wash for 5 min, a linear gradient from 0-100% B over 10 min, followed by a hold at 100% B for 3 min.

### Construction of protein expression plasmids

Genomic DNA from *Variovorax paradoxus* DSM 30034 was isolated using the Wizard Genomic DNA Purification Kit (Promega) following the manufacturer’s instructions. From genomic DNA, *senC* and *egtD* genes were PCR-amplified using Q5 High Fidelity DNA polymerase (NEB) with primers Vpa-SenC-F/R and Vpa-EgtD-F/R, respectively, which have overhangs that allowed for assembly into pET28b(+) (Supplementary Table 3). *SenA* and *senB* from *Variovorax paradoxus* DSM 30034 were obtained as synthetic DNA fragments, codon-optimized for expression in *E. coli* and with overhangs to allow assembly into pET28b(+). Protein expression plasmids were assembled from gene fragments and vector pET28b(+), linearized with NdeI and XhoI (NEB), using HiFi DNA Assembly Master Mix (NEB) following the manufacturer’s instructions. Ligation mixtures were transformed into chemically-competent *E. coli* DH5α by heat-shock and plated onto LB agar containing 50 mg/L kanamycin. After confirmation by Sanger sequencing, assembled plasmids were transformed into *E. coli* BL21(DE3) for protein expression.

### Expression and purification of 6xHis-tagged SenA, SenB, SenC, and EgtD proteins

All four proteins were produced separately in *E. coli* BL21(DE3) cells grown in two 4 L flasks, each containing 2 L Terrific Broth supplemented with 50 mg/L kanamycin at 37°C/170 rpm. Small cultures were prepared by inoculating 40 mL of LB medium containing 50 mg/L Kan with a single colony of *E. coli* BL21(DE3) carrying the desired plasmid. After overnight growth at 37°C/170 rpm, 4 L of TB medium plus 50 mg/L Kan were inoculated with the 40 mL small culture and incubated at 37°C/170 rpm. At OD_600_ = 0.5–0.6, protein expression was induced with 0.2 mM IPTG, and cultures were incubated at 37°C/170 rpm for an additional 12–24 hours. Cells were pelleted by centrifugation (8,000g, 15 min, 4°C), yielding ~7 g of cell paste per L. The cell pastes were stored at −80°C until purification.

All purification steps were carried out in a cold room at 4°C. Cells were resuspended in lysis buffer (5 mL/g cell paste), which consisted of 25 mM Tris-HCl, 300 mM NaCl, 10 mM imidazole, 10% glycerol, pH = 7.7, supplemented with 1 μL/mL Protease Inhibitor Cocktail (Sigma) and 1 mM phenylmethylsulfonyl fluoride. Once homogenous, 0.1 mg/mL deoxyribonuclease I (Alfa Aesar) was added, and the cells were lysed by the addition of 5 mg/mL lysozyme followed by sonication using 30% power (~150 W) in 15 s on/15 s off cycles for a total of 4 min. This process was repeated twice. The lysate was then clarified by centrifugation (17,000g, 15 min, 4°C) and loaded onto a 5 mL Ni-NTA column pre-equilibrated in lysis buffer. The column was washed with lysis buffer and His-tagged proteins were eluted with elution buffer consisting of 25 mM Tris-HCl, 300 mM NaCl, 300 mM imidazole, 10% glycerol, pH = 7.7. Eluted proteins were then buffer-exchanged using a 50 mL column of Sephadex G-25 (Cytiva) into storage buffer consisting of 25 mM Tris-HCl, 150 mM NaCl, 10% glycerol, pH = 7.7. Purified proteins were stored at −80°C. Protein concentrations were determined spectrophotometrically on a Cary 60 UV-visible spectrophotometer (Agilent) using calculated molar extinction coefficients at 280 nm. From 4 L cultures, the following yields were obtained: 105 mg SenA, 198 mg SenB, 152 mg SenC, and 63 mg EgtD.

### In vitro reconstitution of SenC activity

The selenophosphate synthetase activity of SenC was characterized according to previous methods^31–33^. Under anaerobic conditions, a 700 µL reaction containing 2 mM dithiothreitol (DTT), 1 mM ATP, 1.5 mM Na_2_Se, and 20 µM SenC was prepared in buffer consisting of 50 mM tricine, 20 mM KCl, 5 mM MgCl_2_, 10% D_2_O, pH = 7.2. Control reactions were prepared in an identical fashion, lacking either SenC or Na_2_Se, or with Na_2_S in place of Na_2_Se. After a 1-hour incubation period at room temperature, the reactions were transferred to NMR tubes, removed from the glovebox, and immediately analyzed by ^31^P-NMR. Results were consistent with previous reports of selenophosphate synthetase enzymes, with conversion to selenophosphate (SeP) only observed in the presence of all components. No production of thiophosphate was observed in the presence of Na_2_S (Supplementary Fig. 2).

### In vitro reconstitution of SenB activity

Under anaerobic conditions, 200 µL reactions containing 2 mM DTT, 2 mM ATP, 1 mM Na_2_Se, 20 µM SenC, 20 µM SenB, and 2 mM of NDP-sugar were prepared in buffer consisting of 50 mM tricine, 20 mM KCl, 5 mM MgCl_2_, pH = 7.2. Control reactions were prepared in an identical fashion, lacking either SenB, SenC, or ATP. After a 6-hour incubation period at room temperature, reactions were removed from the glovebox and exposed to oxygen for 30 minutes to oxidize any unreacted Na_2_Se. Next, 50 µL of each reaction mixture was quenched with 50 µL of MeOH, while another 50 µL was quenched with 50 µL of 10 mM mBBr in MeCN. Reactions were incubated for an additional 30 min at room temperature in the dark to allow for complete derivatization with mBBr. Reactions quenched with MeOH were filtered and analyzed by LC-MS using a Kinetex Polar C18 column (Phenomenex, 150 × 4.6 mm, 2.6 µm) with a flow rate of 0.4 mL/min and an elution program consisting of 0–20% solvent B over 5 min, followed by 20–100% solvent B over 3 min, and a final step of 3 min at 100%. This assay demonstrated the production of underivatized selenosugar diselenides (Supplementary Fig. 4). Reactions quenched with mBBr were filtered and analyzed by LC-MS using a Synergi Hydro-RP column (Phenomenex, 250 × 4.6 mm, 4 µm) with a flow rate of 1 mL/min and an elution program consisting of a 5% solvent B wash for 3 min, a gradient of 5–75% solvent B over 6 min, followed by a gradient of 75–100% solvent B over 1 min, and a final hold at 100% for 5 min. The remaining 100 µL of each reaction mixture was lyophilized for use as a crude selenosugar substrate in downstream assays with SenA.

### Structural characterization of SenB products

SenB enzymatic assays were carried out anaerobically as described above on a 10-mL scale with UDP-glucose (Glc) and UDP-N-acetylglucosamine (GlcNAc). Reactions were quenched with 3 mL of 10 mM mBBr in MeCN and incubated on a platform rocker for 1 hour in the dark to facilitate complete derivatization. SeGlc-mBBr was purified by semi-preparative HPLC using a Synergi Fusion-RP column (Phenomenex, 250 × 10 mm, 4 µm) with a flow rate of 2.5 mL/min and elution consisting of a gradient of 0–40% solvent B for 15 min, followed by 40–100% solvent B for 3 min, and a final hold at 100% B for 5 min. SeGlcNAc-mBBr was purified by semi-preparative HPLC using a Luna C18 column (Phenomenex, 250 × 10 mm, 5 µm) with a flow rate of 2.5 mL/min and elution consisting of a gradient of 10–50% solvent B for 10 min, followed by a gradient of 50–100% solvent B for 10 min, followed by a hold at 100% for 3 min. Purified compounds were dissolved in DMSO-*d*_*6*_ and analyzed by NMR spectroscopy, confirming the presence of a selenium atom bound to the anomeric carbon, which was determined to be ß-configured for both selenosugars, as shown by the large axial-axial coupling constants between protons on the sugar C-1 and C-2 (9.2 Hz for SeGlc-mBBr and 10.5 Hz for SeGlcNAc-mBBr). Full spectra, chemical shift assignments, and select 2D correlations can be found in Supplementary Figs. 3 and 5 and Supplementary Tables 5 and 6.

### In vitro reconstitution of EgtD activity

Under aerobic conditions, 100 µL reactions containing 0.5 mM histidine, 2 mM S-adenosylmethionine (SAM), and 20 µM EgtD were prepared in buffer consisting of 50 mM Tris-HCl, 150 mM NaCl, pH = 8.0. A control reaction lacking EgtD was prepared in an identical fashion. After a 30-minute incubation period at room temperature, reactions were quenched with 100 µL MeOH, filtered, and analyzed by LC-MS using a Synergi Hydro-RP column (Phenomenex, 250 × 4.6 mm, 4 µm) with a flow rate of 1 mL/min and an elution program gradient consisting of a 0% solvent B wash for 3 min, a followed by a gradient to 100% solvent B over 2 min, and a hold at 100% for 5 min. Results were consistent with previous reports of EgtD enzymes, with conversion of histidine to hercynine observed only in the presence of EgtD^34^ (Supplementary Fig. 7).

### In vitro reconstitution of SenA activity

Under aerobic conditions, crude selenosugar substrates from lyophilized SenBC assay mixtures were first dissolved in buffer consisting of 50 mM Tris-HCl, 150 mM NaCl, pH = 8. Next, 0.5 mM histidine, 2 mM S-adenosylmethionine (SAM), 2 mM DTT, 0.2 mM (NH_4_)_2_Fe(SO_4_)_2_, 20 µM EgtD, and 20 µM SenA were added to a final volume of 100 µL. Control assays were prepared in an identical fashion, except lacking either EgtD or SenA. Assays containing sodium ß-D-thioglucose (Fisher) in place of selenosugar substrate were also prepared in a similar fashion. However, thioglucose was found to be a very poor substrate for SenA, and thus 80 µM enzyme was used to facilitate appreciable conversion. After a 7-hour incubation period at room temperature, 50 µL of each reaction mixture was quenched with 50 µL of MeOH, while another 50 µL was quenched with 50 µL of 10 mM mBBr in MeCN. Reactions were incubated for an additional 30 minutes at room temperature in the dark to allow for complete derivatization with mBBr. The samples were then filtered and analyzed by LC-MS using a Synergi Hydro-RP column (Phenomenex, 250 × 4.6 mm, 4 µm) with a flow rate of 1 mL/min and elution consisting of 5% solvent B for 3 min, a gradient of 5–75% solvent B over 6 min, followed by a gradient of 75–100% over 1 min, and a hold at 100% for 5 min.

### Structural characterization of SenA products

A large-scale SenA enzymatic assay was carried out aerobically as described above using a crude SeGlcNAc substrate from a lyophilized, 10-mL SenBC reaction mixture. Reactions were quenched with 5 mL of 10 mM mBBr in MeCN and incubated on a platform rocker for 1 hour in the dark to facilitate complete derivatization. Selenoneine-mBBr (SEN-mBBr) was purified by semi-preparative HPLC using a Luna C18 column (Phenomenex, 250 × 10 mm, 5 µm) with a flow rate of 2.5 mL/min and an elution program consisting of a gradient of 7.5–80% solvent B for 12 min, followed by a hold at 80% for 4 min. Purified SEN-mBBr was dissolved in D_2_O and analyzed by NMR spectroscopy. The selenium atom was confirmed to be positioned at the imidazole C2 carbon, as evidenced by the absence of an imidazole C-2 proton and diagnostic ^1^H-^13^C HMBC correlations. Full spectra, chemical shift assignments, and select 2D correlations can be found in Supplementary Fig. 8 and Supplementary Table 7. The structures of other SenA products (hercynyl-SeGlcNAc selenoether [GlcNAc-SEN], hercynyl-SeGlc selenoether [Glc-SEN], hercynyl-thioglucose sulfoxide [Glc-EGT=O], and selenoneine [SEN]) were confirmed by tandem HR-MS/MS analysis (Supplementary Fig. 9 and Supplementary Table 8).

## Acknowledgments

We thank Nicole Hauser for helpful discussion as well as the Edward C. Taylor 3^rd^ Year Fellowship in Chemistry (C.M.K.) and the National Science Foundation CAREER Award (M.R.S.) for financial support.

## Author contributions

C.M.K. and M.R.S. conceived the idea for the study. C.M.K. designed and performed the experiments. C.M.K. and M.R.S analyzed data and prepared the manuscript.

## Competing interests

The authors declare that they have no competing interests.

## Additional information

### Supplementary information

This report contains Supplementary Text, Supplementary Tables 1-8, and Supplementary Figs. 1-9.

### Correspondence and requests for materials

should be addressed to Mohammad R. Seyedsayamdost.

## Notes

### Competing Interest Statement

The authors have declared no competing interest.

