## Supplemental file for "A Novel and Prevalent Pathway in Microbial Selenium Metabolism"

**Coding DNA sequences and amino acid sequences of recombinant proteins**

**6xHis-*senA* CDS (from *V. paradoxus*, synthesized, codon optimized)**

atgggcagcagccatcatcatcatcatcacagcagcggcctggtgccgcgcggcagccatatggacagtaccttaccggtgtattcagtt
gctggcgcgcccgaagcgttagcattgcgcgcgggacccccctgcgagtgtgcgggctgcgcttctcgccgcacgccgccgcaccctggat
ctggcggatgattttcgtgcggctctgggagacgcgtatccgggcattggctatgcgccggaactgaacccgccgctgtgggaactgggc
catgtggcgtggtttcaagaatggtggattggccgaaaccgtcagcgcgcacgcggagtggcgtgcgaaccggatcatgcgcgcgaacc
gagcctgctgccgcaagcggatgcttggtatgactccggccgcgtagcgcatcgcacgcgctgggctctgccgctgccggatgcggaagc
gacccgcgattatctggaacgcaccctggcgcagaccctggcgctgctggatgaactgccgccggatgcgcatgatgatgcgctgtatttt
tttcgcctggtggcgctgcatgaagcgatgcatgcggaagcggcggcgtatatggcggaaggcctgggcattgcgctgcgcgaaggcgg
cgtggcgccgcagctggcggaagatgcggaactggaactgccggcgcagcgcctgcgcatgggcagcgatgctgggaccggatttgcg
tttgataacgaactgctgagccatgatgtgagcattgaaccgctgcgcattgatgcgcaagcggtgagctgggcgcgctttctgccgtttg
tggaagcgggcggctatgaacatccggcatggtggtcagatgcgggacgcgattggctggcgcggcaactgctgcgccatcctgcgcat
ttacgggcggctggcaccggctggcagcagagacgtgggggccgttggctgccactggatccgcaaggcgcggcggtgcatttaaatgc
tcacgaagccgaagcgtggtgtcgctgggcgggacgccgactcccgaccgaagcggaatgggaatgcgcggcgctgaccctgccgggc
ttcgcgtggggacgcgtgtgggaatggacgagcagcccgtttgaaccgtatcccggcttcgcgccacacccgtatcgcgattatagcgcg
ccgtggtttggcacccgcagagtgctgcgcggcgcgtgtcatgcgacgagcgcggcgctggcccacgcgcgctatcgcaactttttgaa
ccgcatcgccgcgatatttttgcgggctttcgcagctgccgcgcgccgggcggctaa

**6xHis-SenA protein sequence**

MGSSHHHHHHSSGLVPRGSHMDSTLPVYSVAGAPEALALRAGPPASVRAALLAARRRTLDLADDFRAALGD
AYPGIGYAPELNPPLWELGHVAWFQEWWIGRNRQRARGVACEPDHAREPSLLPQADAWYDSGRVAHRT
RWALPLPDAEATRDYLERTLAQTLALLDELPPDAHDDALYFFRLVALHEAMHAEAAAYMAEGLGIALREGGV
APQLAEDAELELPAQRLRMGSDAGTGFAFDNELLSHDVSIEPLRIDAQAVSWARFLPFVEAGGYEHPAWWS
DAGRDWLARQLLRHPAHLRAAGTGWQQRRGGRWLPLDPQGAAVHLNAHEAEAWCRWAGRRLPTEAE
WECAALTLPGFAWGRVWEWTSSPFEPYPGFAPHPYRDYSAPWFGTRRVLRGACHATSAALAHARYRNFFE
PHRRDIFAGFRSCRAPGG*

**6xHis-*senB* CDS (from *V. paradoxus*, synthesized, codon optimized)**

atgggcagcagccatcatcatcatcatcacagcagcggcctggtgccgcgcggcagccatatgagcaacccgagcctggtgattgtgag
cccggcgctgccgggcgcgaacaacggcaactggcgcaccgctcaacgctggaaagcgctgctgagccctgtgtgctccgcacgcgttg
tgcagcagtggccggatgcggatgcgagcgcggataccgtgatgctggcgctgcatgcgcgccgcagcgcggaaagcattgcgcattgg
gcgcatgcgcatccgggccgcggcctgggcgtggtgctgaccggcaccgatctgtatcaagatatcggtagcgaccctcaagcgcagag
gtcactgcagctggcgcagcgactggtggtgttgcaagcgctgggcgcggaagctttacccccggaatgccgcgcgaaagcccgcgtgg
tgtatcagagcacgagcgcgcgtgcggaactgccgaaaagcgcgcgtcagctgcgcgcggtgatggtgggccatctgcgccaagtgaa
aagcccgcagaccctgtttgatgcggcgcgcctgctgtgcggccgcgaagatattcgcattgatcatattggggacgcgggcgatgccgg
ccttggagaactcgcgcgtgcgcttgcgtcagattgccctgggtacagatggttgggcgcgctgccgcatgcgcagacccgtcagcgcat
tcagcgcgcgcatgtgctggtgcatacgagcgcgctggaaggcggcgcgcatgtgattatggaagcggtgcgcagcggcaccccggtgc
tggcgagccgcgtgccgggcaacgtgggcatgctgggcaacgattatgcgggctattttccgcatggtgatgcggcggcactggcggca
ctgctggaggcgtgccgcgcgggccaagggagcaaagatcgcgcagcgggcttactggattcgctgcgcacgcagtgcgcgctccgtgc
gccgctgtttgatccgcgcgcggaacaagcggcgctgtttcagctgctgaacgaactgcagccgccgccgccgtaa

**6xHis-SenB protein sequence**

MGSSHHHHHHSSGLVPRGSHMSNPSLVIVSPALPGANNGNWRTAQRWKALLSPVCSARVVQQWPDADA
SADTVMLALHARRSAESIAHWAHAHPGRGLGVVLTGTDLYQDIGSDPQAQRSLQLAQRLVVLQALGAEALP

PECRAKARVVYQSTSARAELPKSARQLRAVMVGHLRQVKSPQTLFDAARLLCGREDIRIDHIGDAGDAGLGE
LARALASDCPGYRWLGALPHAQTRQRIQRAHVLVHTSALEGGAHVIMEAVRSGTPVLASRVPGNVGMLGN
DYAGYFPHGDAAALAALLEACRAGQGSKDRAAGLLDSLRTQCALRAPLFDPRAEQAALFQLLNELQPPPP*

**6xHis-*senC* CDS (from *V. paradoxus*, cloned from genomic DNA)**
atgggcagcagccatcatcatcatcatcacagcagcggcctggtgccgcgcggcagccatatgaacgctgccctcccgcaagcccccgc
cgccgagccccgcctcacctcgctgtcgcacggcggcggctgcggctgcaagatcgcgcccggcgtgctgtcggaaatcctcaagggca
cggcgcgcatgccgatgccgcccgagctgctggtcggcatcgagacggccgacgacgcggccgtctaccagctcaacgacgagcaggc
gctgatcgccaccaccgacttcttcatgccgatcgtcgacgatccgttcgacttcggccgcatcgcggccaccaacgcgatctccgacgtg
tatgcgatgggcggcaagccgatcctggcgctcgcgctggtcggcatgccgatcaacgtgctcagcacggcgaccatcgggcgcatcctc
gaaggcggcgaatcggtgtgccgggcggcgggcattccgatcgcgggcggccacaccatcgattcggtcgaggccatctacgggctggt
cgcgctcggcctggtccacccgaagcgcgtcaagcgcaatgccggcgccaaggccggcgacctgctggtgctgggcaagccgctgggc
gtgggcatcctgtcggcggcgctcaagaaggaggcactcgacgcggccggctaccaacgcatgatcgccaccacgacgcggctgaaca
cgtccggccccgagctcgcggcgctcgacggcgtgcatgcgctgaccgacgtgacgggcttcggcttggccggccatgcgctggaactg
gcacgcggcgcaaagctgtcgctggagatcgactggcagcaggtgcccgtgctggaaggcgtgcgcggcctggtgcagtcgggacacg
tcaccggcgcatccggccgcaactgggccggctacggctccgatgtggcgctgccgcagaactttgccgaggaagagcgcgccctgctc
accgacccgcagaccagcggcggactgctggtttcgtgcgcgcccgaggcgctcgacgaggtgctggccatcttccgccggcatggcttc
gacgacgccgcggtgattggcactgccagcgcgcaagggagcgcgccgcggctggtgatccgctga

**6xHis-SenC protein sequence**
MGSSHHHHHHSSGLVPRGSHMNAALPQAPAAEPRLTSLSHGGGCGCKIAPGVLSEILKGTARMPMPPELL
VGIETADDAAVYQLNDEQALIATTDFFMPIVDDPFDFGRIAATNAISDVYAMGGKPILALALVGMPINVLSTA
TIGRILEGGESVCRAAGIPIAGGHTIDSVEAIYGLVALGLVHPKRVKRNAGAKAGDLLVLGKPLGVGILSAALKK
EALDAAGYQRMIATTTRLNTSGPELAALDGVHALTDVTGFGLAGHALELARGAKLSLEIDWQQVPVLEGVR
GLVQSGHVTGASGRNWAGYGSDVALPQNFAEEERALLTDPQTSGGLLVSCAPEALDEVLAIFRRHGFDDAA
VIGTASAQGSAPRLVIR*

**6xHis-*egtD* CDS (from *V. paradoxus*, cloned from genomic DNA)**
atgggcagcagccatcatcatcatcatcacagcagcggcctggtgccgcgcggcagccatatgcgcaatccaagctcctcgtcctcctttt
cgtttgccgccgcccagccgcagcagcgccagcctgtcgcgcctgccgccgcggaaaaaaatgcggcgctgctctccgagttcggccgc
gagatgcaggcgggcctcgcgcgcaggccgcgcagcatttcgcccaagtatttttacgacgccgcgggttcgcagctgttcgaccgcatc
tgcgagctgcccgagtactaccccacgcgcaccgagctgcgcattcttggcgaatgcgcgggcgagatcgccgcgcaggtgggtccggg
cgccgagatcgtcgagttcggcgccggctcgctcaccaaggtgcggctgctgctcgatgcgctcgaggcgcccaggcgctacctgccgat
cgacatttccggcgagcacctggcggccgccgccgagcggctcagggccgactacccgcagctggcggtgcagccgatcgcggcggac
tacaccatgccgctggtgctgcccgcgccgctcgccggtgcgggcaagcgcgtgggtttttccccggctcgacgatcggaaatttcgagc
ccgacgaggcactggccttcctgcagctcgcggcgcgcatgctgcgcggcggcgggctgctgatcggcgtcgacctggtgaaggacccg
ggccgcctgcatgcggcctacaacgacgcgcagggcgtgaccgcggcgttcaacctcaacctgctgcggcgcgccaacgccgagctcga
cgcgaacttcgatctcgagggctttgcgcacgcggcgttctacaacgcgccgaaacagcgcatcgagatgcacctggtgagccggcggg
accagatcgtttcgctgaacggcgagcggtttgcgttcgaggaaggggagaccattcacaccgagtactcgcacaagttcactgtggaa
ggattgcgggcgctggctgtgaaggcggggttccggccgggggctgtgtggacggatccggagaggctcttcagcgtgcattggttgggg
gcttga

**6xHis-EgtD protein sequence**

MGSSHHHHHHSSGLVPRGSHMRNPSSSSSFSFAAAQPQQRQPVAPAAAEKNAALLSEFGREMQAGLARR
PRSISPKYFYDAAGSQLFDRICELPEYYPTRTELRILGECAGEIAAQVGPGAEIVEFGAGSLTKVRLLLDALEAPR
RYLPIDISGEHLAAAAERLRADYPQLAVQPIAADYTMPLVLPAPLAGAGKRVGFFPGSTIGNFEPDEALAFLQL
AARMLRGGGLLIGVDLVKDPGRLHAAYNDAQGVTAAFNLNLLRRANAELDANFDLEGFAHAAFYNAPKQRI
EMHLVSRRDQIVSLNGERFAFEEGETIHTEYSHKFTVEGLRALAVKAGFRPGAVWTDPERLFSVHWLGA*

**Supplementary Table 1.** Protein families of the ten most common *selD*-overlapping genes.

| Family | # overlapping | % of overlapping genes | % of all SelDs |
|---|---|---|---|
| SelU | 1302 | 44 | 12 |
| TIGR04348 | 143 | 4.8 | 1.3 |
| YedF | 131 | 4.4 | 1.2 |
| SelA | 86 | 2.9 | 0.8 |
| DUF3343 | 45 | 1.5 | 0.41 |
| GalE | 37 | 1.3 | 0.34 |
| Sec-lyase | 37 | 1.3 | 0.34 |
| LysR | 34 | 1.1 | 0.31 |
| NrfD | 34 | 1.1 | 0.31 |
| SelB | 32 | 1.1 | 0.29 |

**Supplementary Table 2.** Bacterial strains used or generated in this study.

| Strain | Purpose | Source |
|---|---|---|
| *Amycolatopsis palatopharyngis* DSM 44832 | Actinomycete containing *sen* and *egt* clusters, production screening for SEN and EGT | DSMZ |
| *Variovorax paradoxus* DSM 30034 | Betaproteobacterium containing *sen* and *egt* clusters, production screening for SEN and EGT | DSMZ |
| *Streptomyces rimosus* ATCC 10970 | Actinomycete containing *egt* cluster only, production screening for SEN and EGT | ATCC |
| *Burkholderia thailandensis* E264 | Betaproteobacterium containing *egt* cluster only, production screening for SEN and EGT | ATCC |
| *E. coli* DH5α | Host strain for cloning | NEB |
| *E. coli* BL21(DE3) | Host strain for protein expression | NEB |
| *E. coli* BL21(DE3) + pET-28b(+)_6xHis-SenA | Host strain for expression of 6xHis-SenA | This study |
| *E. coli* BL21(DE3) + pET-28b(+)_6xHis-SenB | Host strain for expression of 6xHis-SenB | This study |
| *E. coli* BL21(DE3) + pET-28b(+)_6xHis-SenC | Host strain for expression of 6xHis-SenC | This study |
| *E. coli* BL21(DE3) + pET-28b(+)_6xHis-EgtD | Host strain for expression of 6xHis-EgtD | This study |

**Supplementary Table 3.** Primers used in this work.

| Primer | Sequence (5'->3') | Purpose |
|---|---|---|
| Vpa-SenC-F | tggtgccgcgcggcagccatatgaacgctgccctccc | Amplification of *senC* for assembly into pET-28b(+) |
| Vpa-SenC-R | agtggtggtggtggtggtgctcagcggatcaccagcc | |
| Vpa-EgtD-F | tggtgccgcgcggcagccatatgcgcaatccaagctcc | Amplification of *egtD* for assembly into pET-28b(+) |
| Vpa-EgtD-R | agtggtggtggtggtggtgctcaagcccccaaccaatg | |

**Supplementary Table 4.** Plasmids used or generated in this study.

| Plasmid | Purpose | Source |
|---|---|---|
| pET-28b(+) | KanR, Expression vector | Novagen |
| pET-28b(+)_6xHis-SenA | Expression of 6xHis-SenA | This study |
| pET-28b(+)_6xHis-SenB | Expression of 6xHis-SenB | This study |
| pET-28b(+)_6xHis-SenC | Expression of 6xHis-SenC | This study |
| pET-28b(+)_6xHis-EgtD | Expression of 6xHis-EgtD | This study |

**Supplementary Table 5**. NMR assignments for SeGlc-mBBr. The structure and numbering scheme for SeGlc-mBBr are shown at the top of the table. See Supplementary Fig. 3 for NMR spectra.

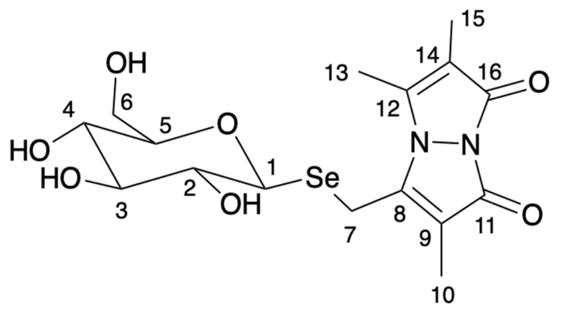


| Pos | δH (m, J in Hz) | δC |
|---|---|---|
| 1 | 4.75 (d, J = 9.2 Hz) | 80.1 |
| 2 | 3.20 (m) | 78.3 |
| 3 | 3.18 (m) | 74.4 |
| 4 | 3.16 (m) | 70.3 |
| 5 | 3.19 (m) | 82.7 |
| 6 | 3.53 (dd, J = 12.2, 6.0 Hz)<br>3.75 (dd, J = 12.2, 1.8 Hz) | 61.6 |
| 7 | 4.00 (d, J = 13.4 Hz)<br>4.17 (d, J = 13.4 Hz) | 12.3 |
| 8 | - | 149.6 |
| 9 | - | 111.9 |
| 10 | 1.83 (s) | 7.0 |
| 11 | - | 160.3 |
| 12 | - | 147.9 |
| 13 | 2.48 (d, J = 0.9 Hz) | 12.0 |
| 14 | - | 111.0 |
| 15 | 1.80 (d, J = 0.9 Hz) | 6.8 |
| 16 | - | 160.6 |

**Supplementary Table 6.** NMR assignments for SeGlcNAc-mBBr. The structure and numbering scheme for SeGlcNAc-mBBr are shown at the top of the table. Key NMR correlations for solving the structure are shown below the table with thick lines and red arrows indicating COSY and HMBC correlations, respectively. See Supplementary Fig. 5 for NMR spectra.

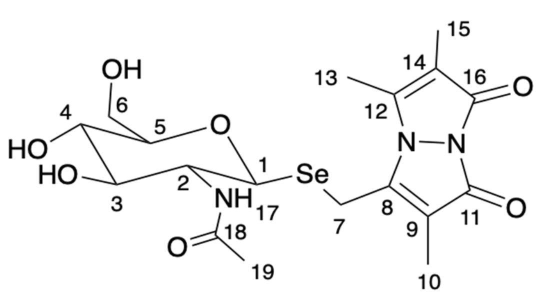


| Pos | δH (m, J in Hz) | δC |
|---|---|---|
| 1 | 4.61 (d, 10.5) | 80.1 |
| 2 | 3.56 (dd, 9.8, 9.2) | 55.2 |
| 3 | 3.19 (dd, 9.8, 7.9) | 75.4 |
| 4 | 3.05 (m) | 70.8 |
| 5 | 3.03 (m) | 82.8 |
| 6 | 3.40 (dd, 11.9, 5.5)<br>3.63 (dd, 12.0, 1.7) | 61.6 |
| 7 | 3.83 (d, 13.5)<br>4.01 (d, 13.4) | 13.0 |
| 8 | - | 149.2 |
| 9 | - | 112.1 |
| 10 | 1.68 (s) | 7.0 |
| 11 | - | 160.3 |
| 12 | - | 147.9 |
| 13 | 2.32 (s) | 11.9 |
| 14 | - | 111.0 |
| 15 | 1.65 (s) | 6.7 |
| 16 | - | 160.6 |
| 17 | 7.80 (d, 9.2) | - |
| 18 | - | 169.9 |
| 19 | 1.70 (s) | 23.2 |

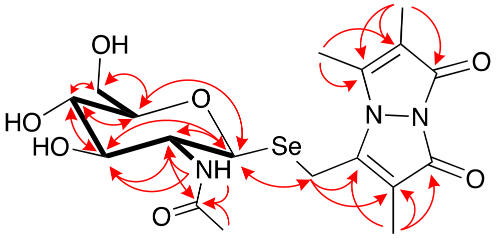

**Supplementary Table 7.** NMR assignments for SEN-mBBr The structure and numbering scheme for SEN-mBBr are shown at the top of the table. See Supplementary Fig. 8 for NMR spectra.

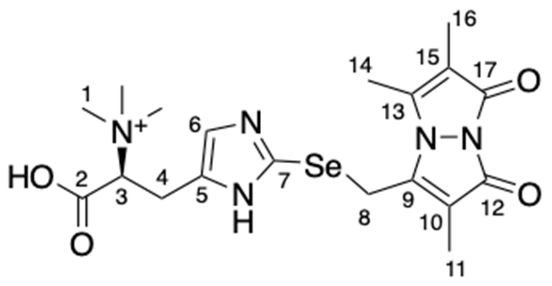


| Pos | δH (m) | δC |
|---|---|---|
| 1 | 3.14 (s) | 52.0 |
| 2 | - | 170.4 |
| 3 | 3.69 (m) | 78.0 |
| 4 | 3.07 (m) | 25.0 |
| 5 | - | 135.5 |
| 6 | 6.97 (s) | 120.6 |
| 7 | - | 128.1 |
| 8 | 3.97 (s) | 19.0 |
| 9 | - | 148.5 |
| 10 | - | 112.7 |
| 11 | 1.22 (s) | 4.8 |
| 12 | - | 162.2 |
| 13 | - | 149.5 |
| 14 | 2.29 (s) | 11.0 |
| 15 | - | 111.3 |
| 16 | 1.68 (s) | 5.4 |
| 17 | - | 162.9 |

**Supplementary Table 8.** Fragments observed by HR-MS/MS analysis of SenA reaction mixtures. Predicted species correspond to fragment ions observed for each of the four parent ions analyzed. Ion numbers correspond to labels in the HR-MS/MS profiles shown in Supplementary Fig. 9.

| Ion # | Predicted Species | m/z calc. | Parent ion | | | | | | | |
|---|---|---|---|---|---|---|---|---|---|---|
| | | | GlcNAc-SEN | | Glc-SEN | | SEN | | Glc-EGT=O | |
| | | | m/z obs. | error (ppm) | m/z obs. | error (ppm) | m/z obs. | error (ppm) | m/z obs. | error (ppm) |
| 1 | 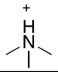 | 60.0808 | 60.0806 | -3.3 | 60.0806 | -3.3 | 60.0807 | -1.7 | 60.0814 | 10.0 |
| 2 | 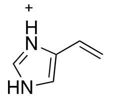 | 95.0604 | 95.0599 | -5.3 | 95.0600 | -4.2 | 95.0602 | -2.1 | n/a | n/a |
| 3 | 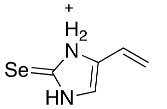 | 174.9769 | 174.9760 | -5.1 | 174.9758 | -6.3 | 174.9761 | -4.6 | n/a | n/a |
| 4 | 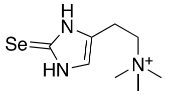 | 234.0504 | 234.0497 | -3.0 | 234.0493 | -4.7 | 234.0498 | -2.6 | n/a | n/a |
| 5 | 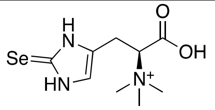 | 278.0402 | 278.0394 | -2.9 | 278.0389 | -4.7 | 278.0390 | -4.3 | n/a | n/a |
| 6 | 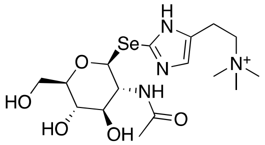 | 437.1298 | 437.1303 | 1.1 | n/a | n/a | n/a | n/a | n/a | n/a |
| 7 | 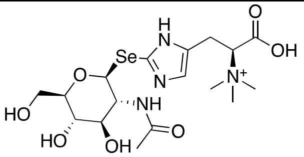 | 481.1196 | 481.1185 | -2.3 | n/a | n/a | n/a | n/a | n/a | n/a |
| 8 | 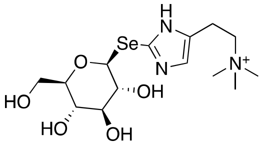 | 396.1032 | n/a | n/a | 396.1016 | -4.0 | n/a | n/a | n/a | n/a |
| 9 | 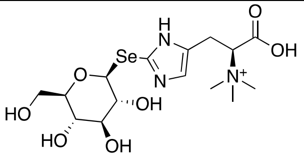 | 440.093 | n/a | n/a | 440.0916 | -3.2 | n/a | n/a | n/a | n/a |
| 10 | 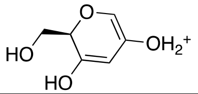 | 145.0495 | n/a | n/a | n/a | n/a | n/a | n/a | 145.0506 | 7.6 |
| 11 | 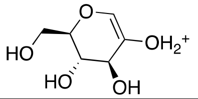 | 163.0601 | n/a | n/a | n/a | n/a | n/a | n/a | 163.0607 | 3.7 |

| | | | | | | | | | | |
|---|---|---|---|---|---|---|---|---|---|---|
| 12 | 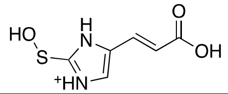 | 187.0172 | n/a | n/a | n/a | n/a | n/a | n/a | 187.0177 | 2.7 |
| 13 | 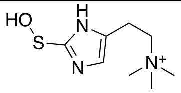 | 202.1009 | n/a | n/a | n/a | n/a | n/a | n/a | 202.1013 | 2.0 |
| 14 | 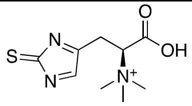 | 228.0801 | n/a | n/a | n/a | n/a | n/a | n/a | 228.0804 | 1.3 |
| 15 | 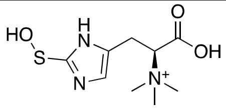 | 246.0907 | n/a | n/a | n/a | n/a | n/a | n/a | 246.0915 | 3.3 |
| 16 | 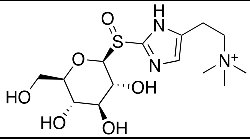 | 364.1537 | n/a | n/a | n/a | n/a | n/a | n/a | 364.1533 | -1.1 |
| 17 | 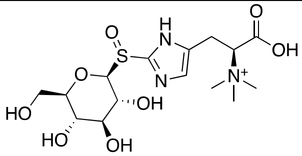 | 408.1435 | n/a | n/a | n/a | n/a | n/a | n/a | 408.1435 | 0.0 |

**Supplementary Figure 1.** Representative *sen* gene clusters in selected bacteria. The top two strains were examined in this work and shown to produce selenoneine.

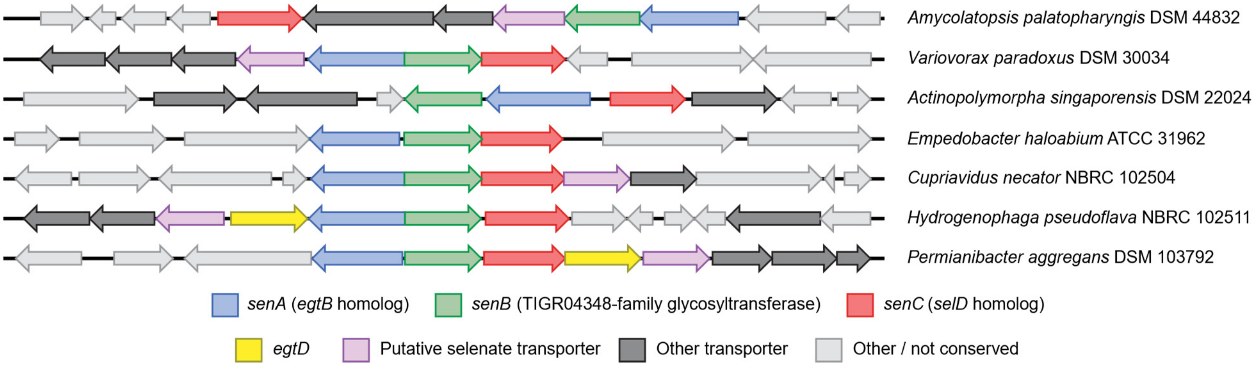

**Supplementary Figure 2.** Monitoring SenC activity by $^{31}$P-NMR. Reaction content for each spectrum is shown on the left. The peaks are labeled. SeP was only observed when all components (SenC, $Na_2Se$, and ATP) were present.

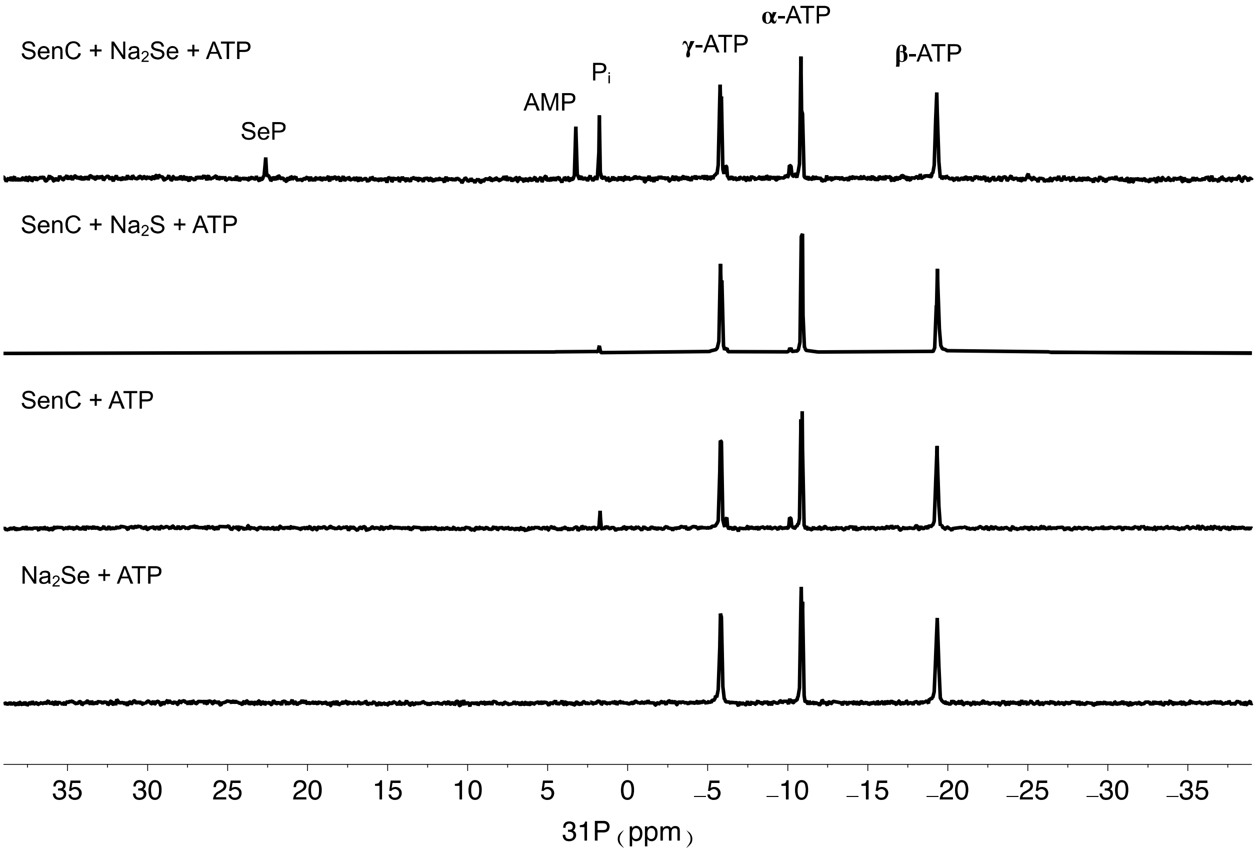

**Supplementary Figure 3.** NMR spectra of SeGlc-mBBr in DMSO-$d_6$ at 500 MHz on pages S13–S16. Shown are from top to bottom $^1$H, $^1$H-$^1$H COSY, $^1$H-$^{13}$C HSQC, and $^1$H-$^{13}$C HMBC spectra.

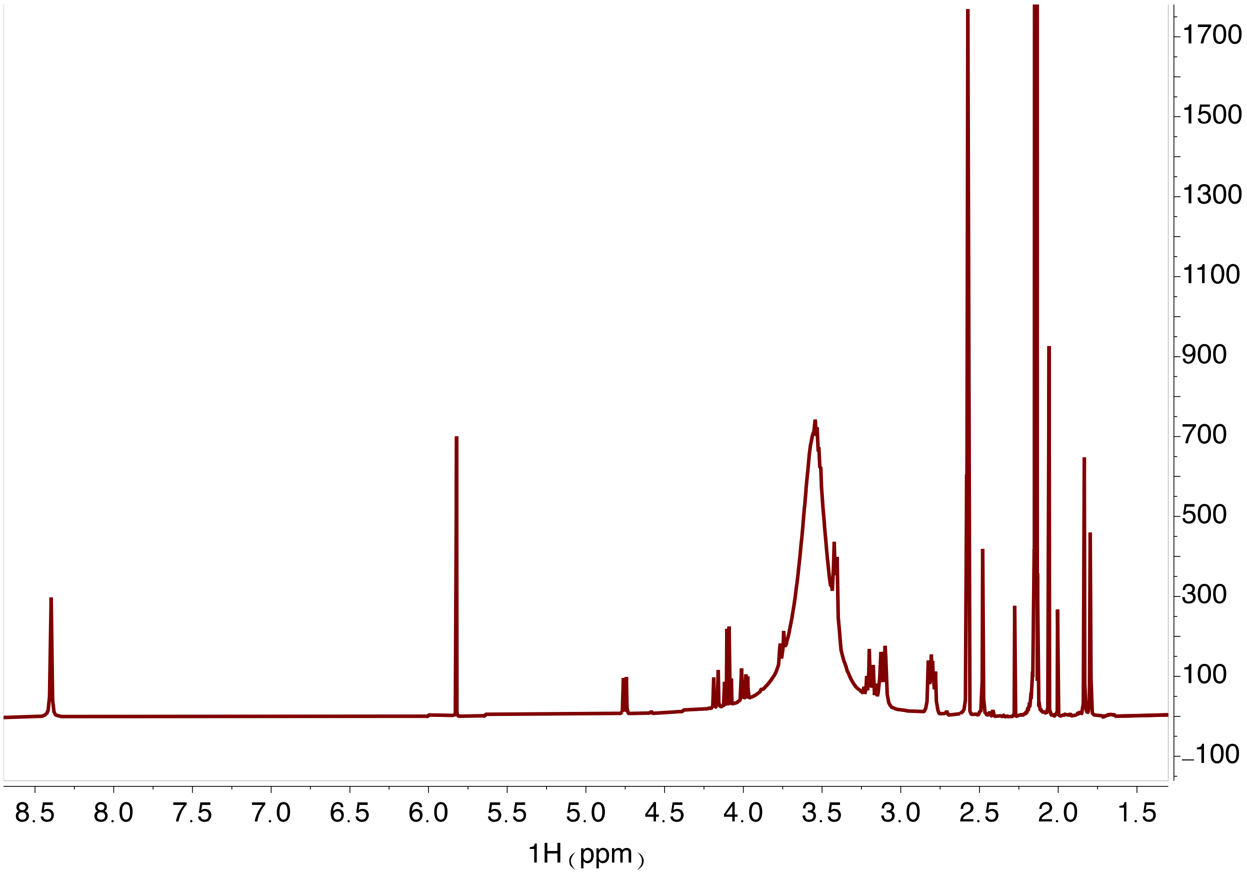

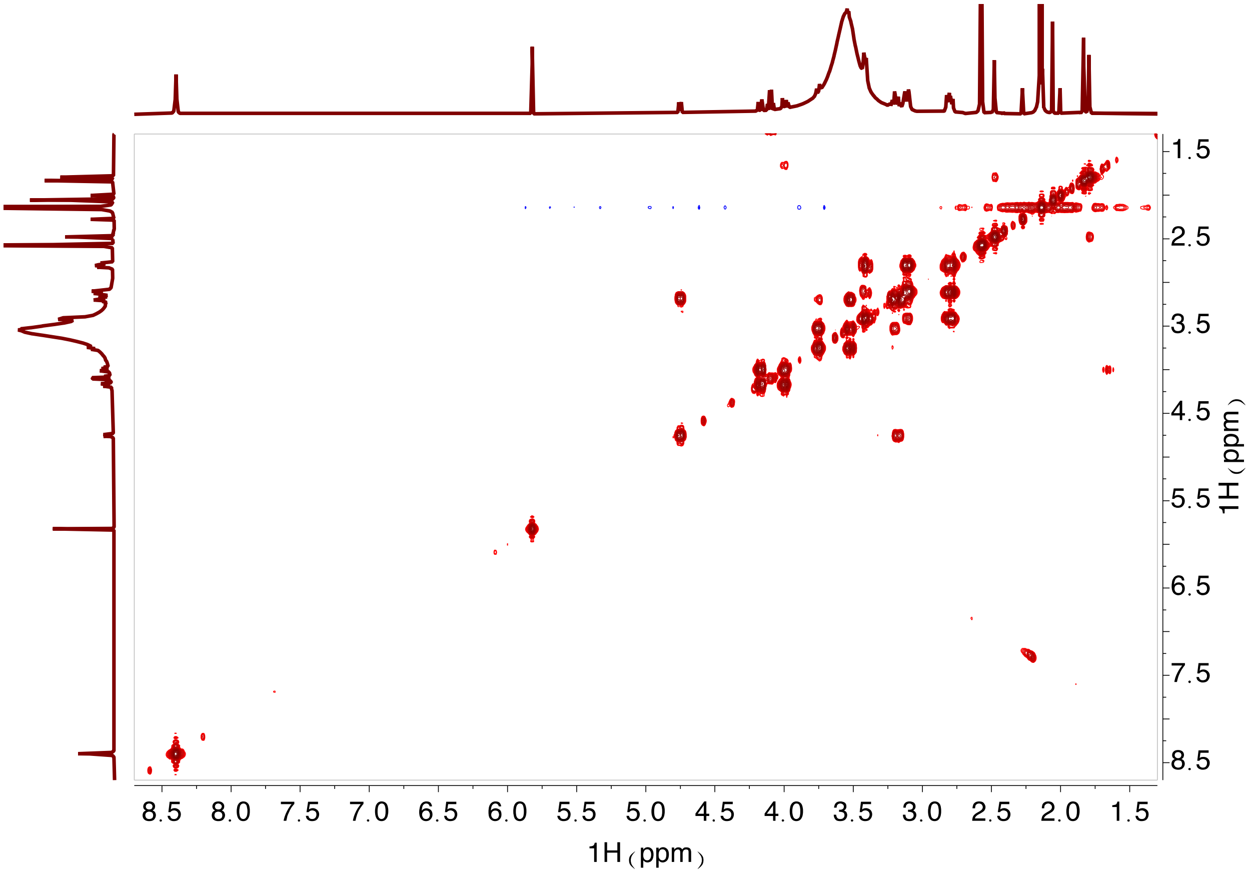

8.5
8.0
7.5
7.0
6.5
6.0
5.5
5.0
4.5
4.0
3.5
3.0
2.5
2.0
1.5
1H（ppm）
1.5
2.5
3.5
4.5
5.5
6.5
7.5
8.5
1H（ppm）

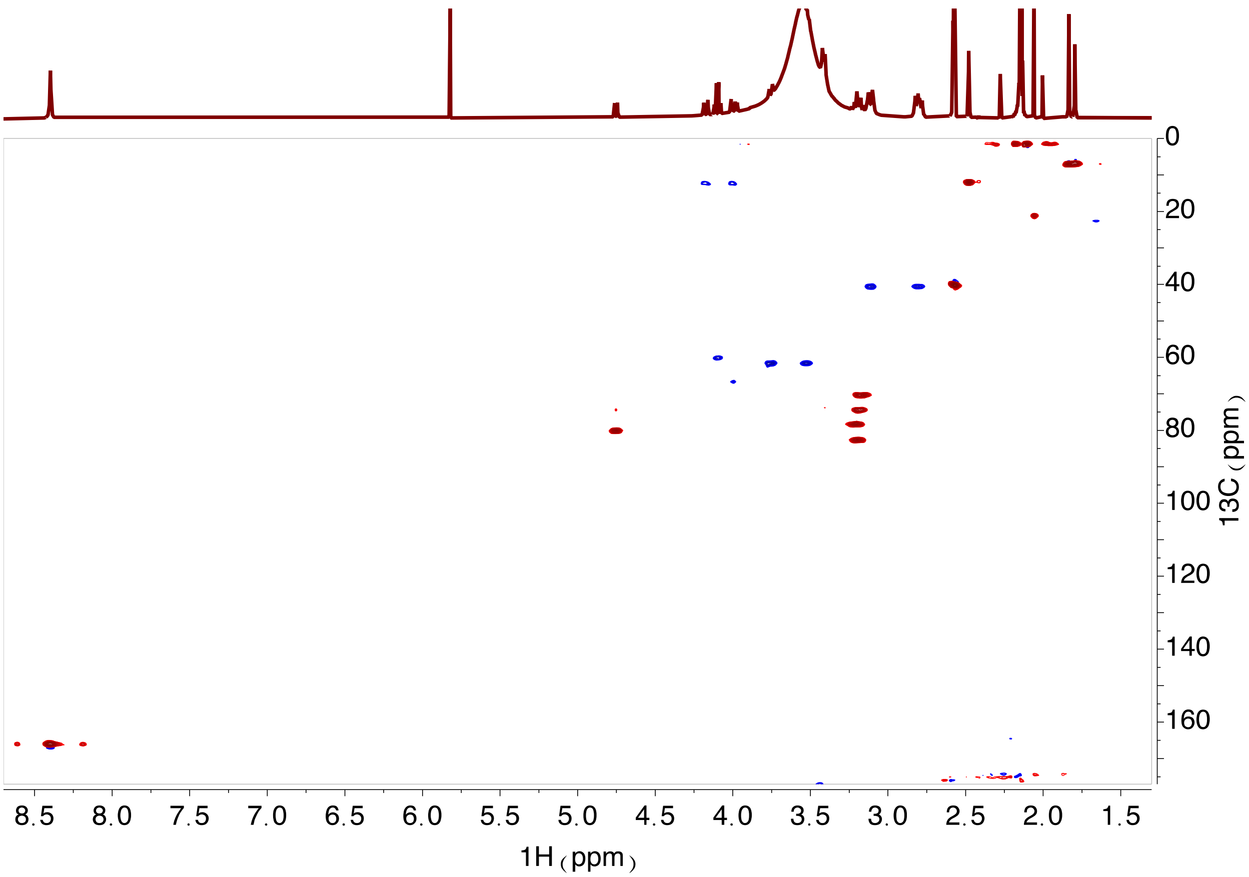
0
20
40
60
80
100
120
140
160
13C（ppm）
8.5 8.0 7.5 7.0 6.5 6.0 5.5 5.0 4.5 4.0 3.5 3.0 2.5 2.0 1.5
1H（ppm）

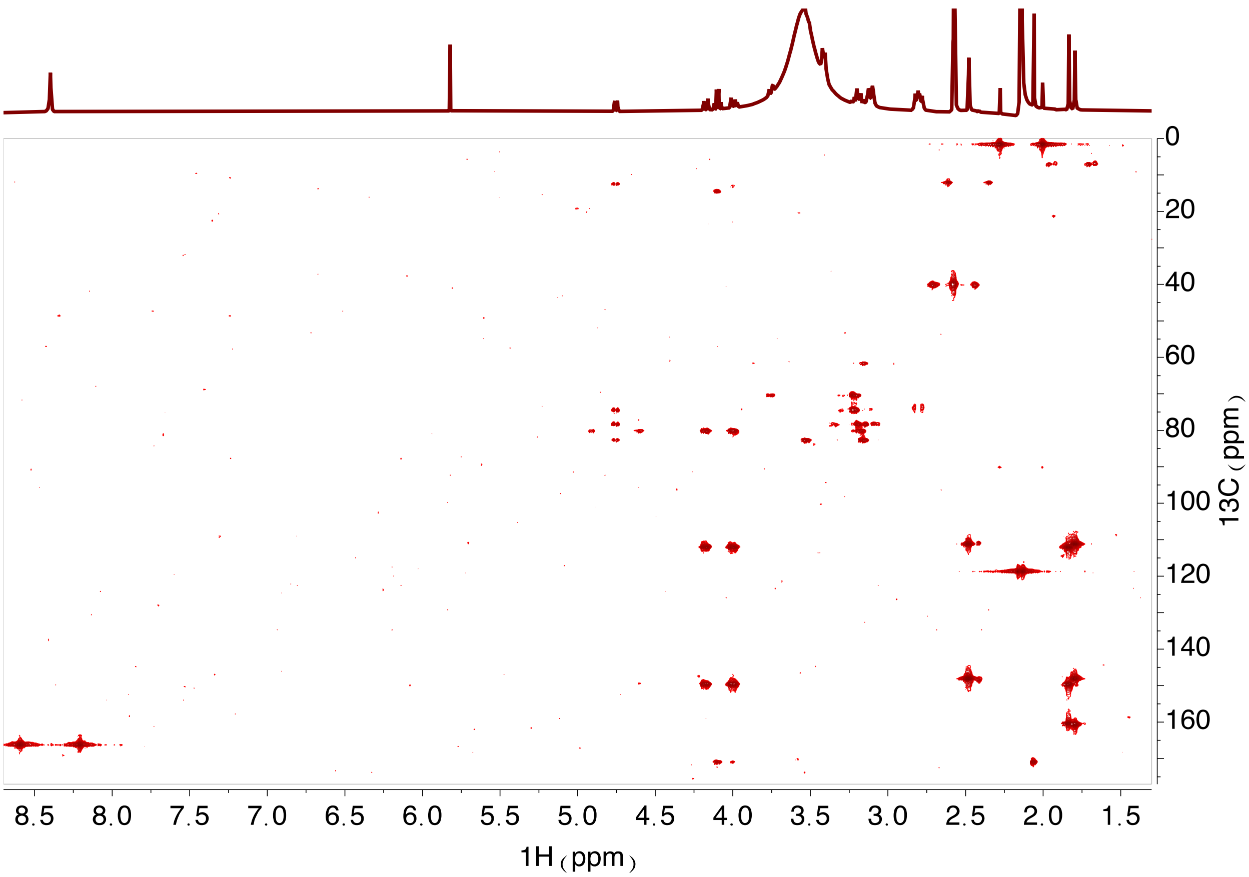

1H（ppm）
13C（ppm）
8.5 8.0 7.5 7.0 6.5 6.0 5.5 5.0 4.5 4.0 3.5 3.0 2.5 2.0 1.5
0 20 40 60 80 100 120 140 160

**Supplementary Figure 4.** Extracted ion chromatograms (panels **a**, **c**, and **e**) and HR-MS profiles (panels **b**, **d**, and **f**) of selenosugar diselenides from underivatized SenBC reaction mixtures.

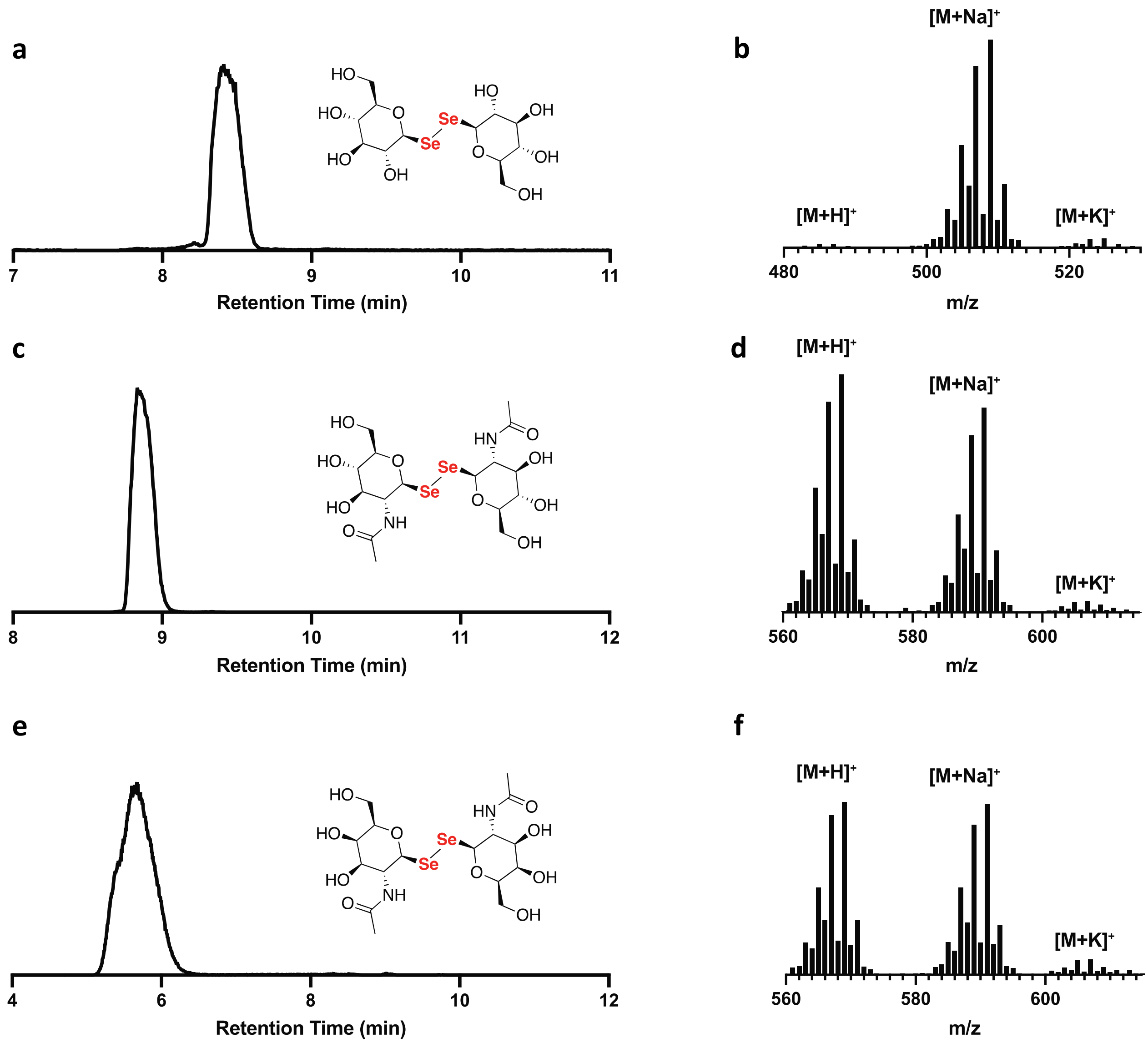

**Supplementary Figure 5.** NMR spectra of SeGlcNAc-mBBr in DMSO-$d_6$ at 500 MHz on pages S18–S21. Shown are from top to bottom $^1$H, $^1$H-$^1$H COSY, $^1$H-$^{13}$C HSQC, and $^1$H-$^{13}$C HMBC spectra.

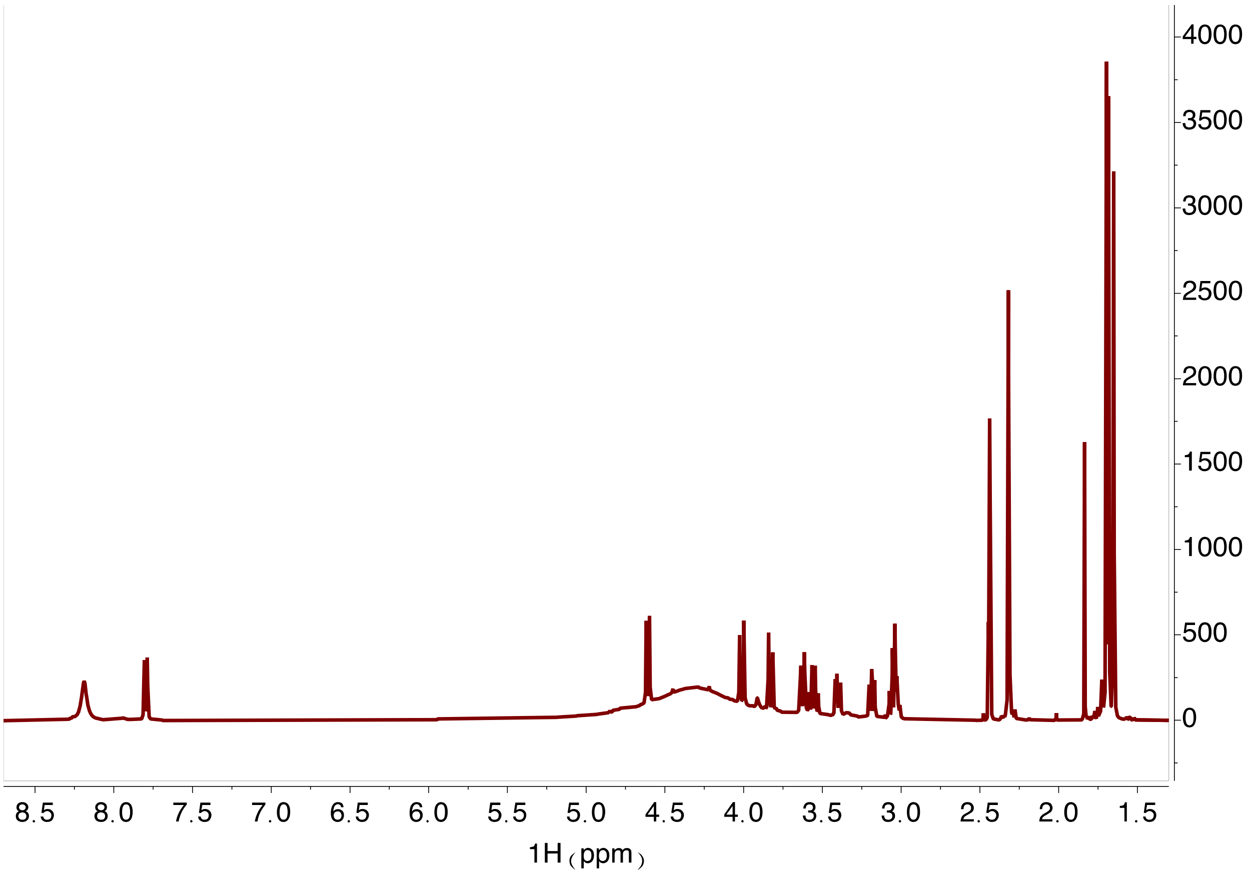

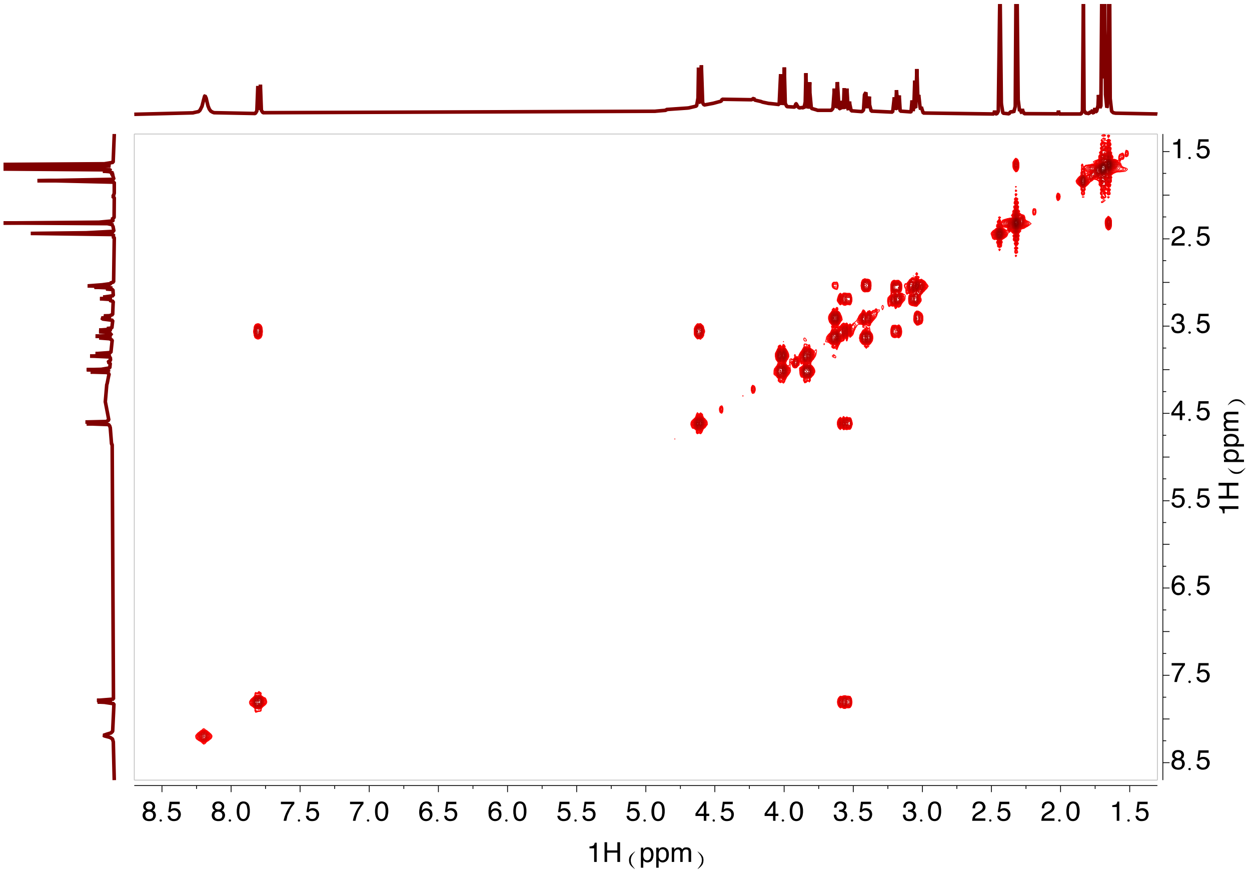

8.5 8.0 7.5 7.0 6.5 6.0 5.5 5.0 4.5 4.0 3.5 3.0 2.5 2.0 1.5
1H（ppm）
1.5 2.5 3.5 4.5 5.5 6.5 7.5 8.5
1H（ppm）

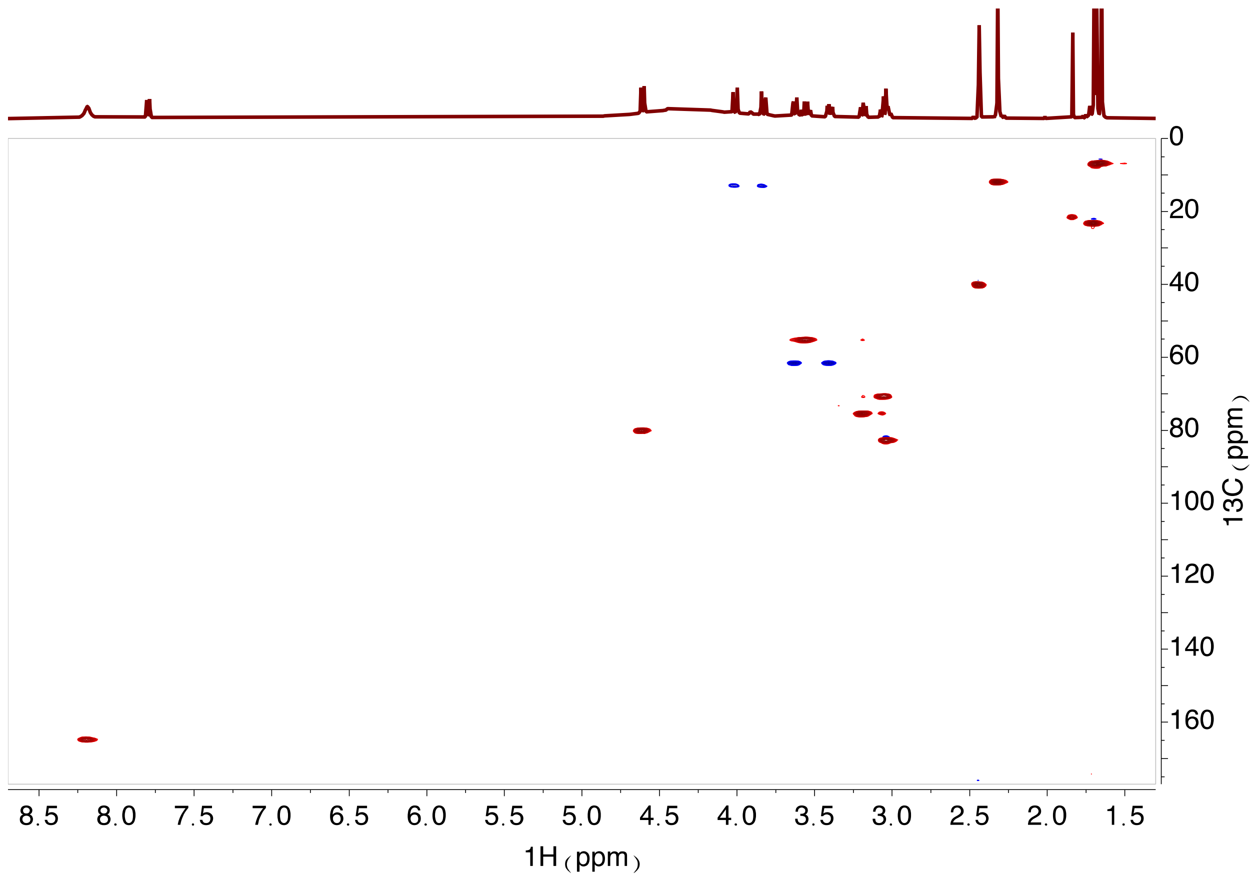

0
20
40
60
80
100
120
140
160
13C（ppm）
8.5
8.0
7.5
7.0
6.5
6.0
5.5
5.0
4.5
4.0
3.5
3.0
2.5
2.0
1.5
1H（ppm）

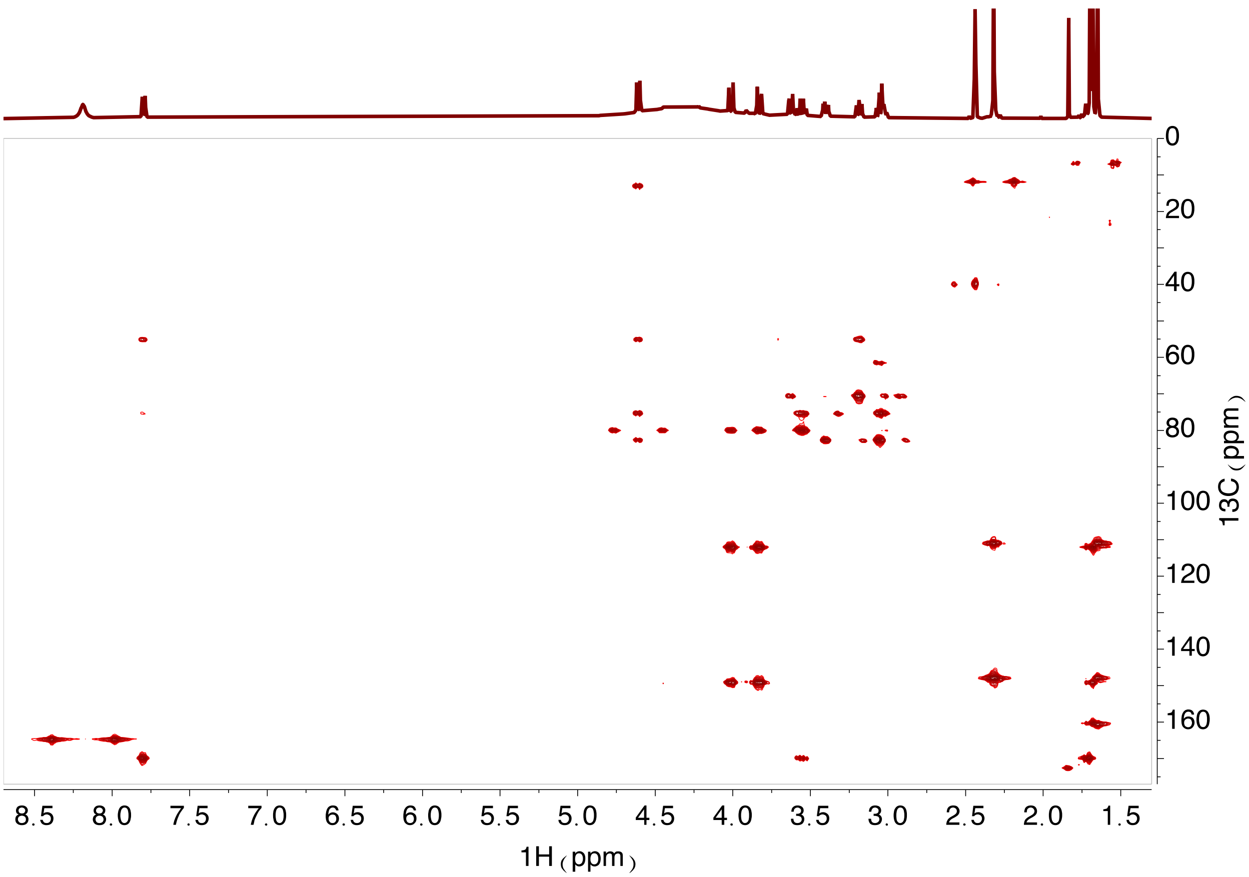

0
20
40
60
80
100
120
140
160
13C (ppm)
8.5
8.0
7.5
7.0
6.5
6.0
5.5
5.0
4.5
4.0
3.5
3.0
2.5
2.0
1.5
1H (ppm)

**Figure S6.** Logo plots displaying multiple sequence alignments of SenA and EgtB proteins. Amino acids are numbered with respect to the structurally characterized EgtB from *M. thermoresistibile*. The catalytic tyrosine and the iron-binding residues are conserved. Some divergence is observed in the thiol- and hercynine-binding residues.

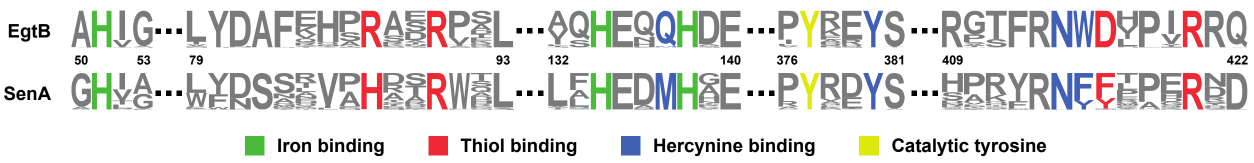


**Supplementary Figure 7.** Reconstitution of EgtD activity in vitro. Shown is the reaction catalyzed by *V. paradoxus* EgtD (**a**) and detection of the product (blue peak) by HPLC-MS (**b**).

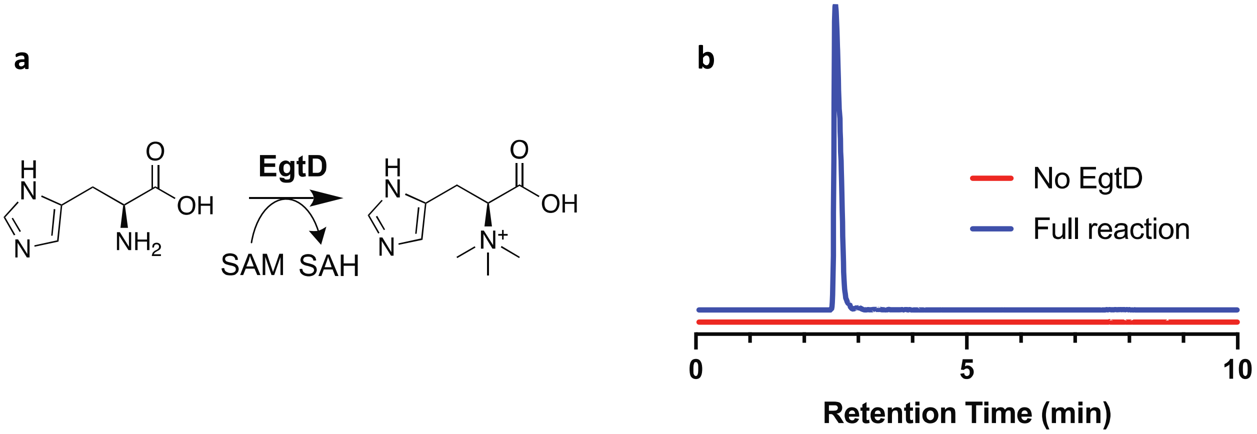

**Supplementary Figure 8.** NMR spectra of selenoneine-mBBr in $D_2O$ at 500 MHz on pages S23–S26. Shown are from top to bottom $^1H$, $^1H$-$^1H$ COSY, $^1H$-$^{13}C$ HSQC, and $^1H$-$^{13}C$ HMBC spectra.

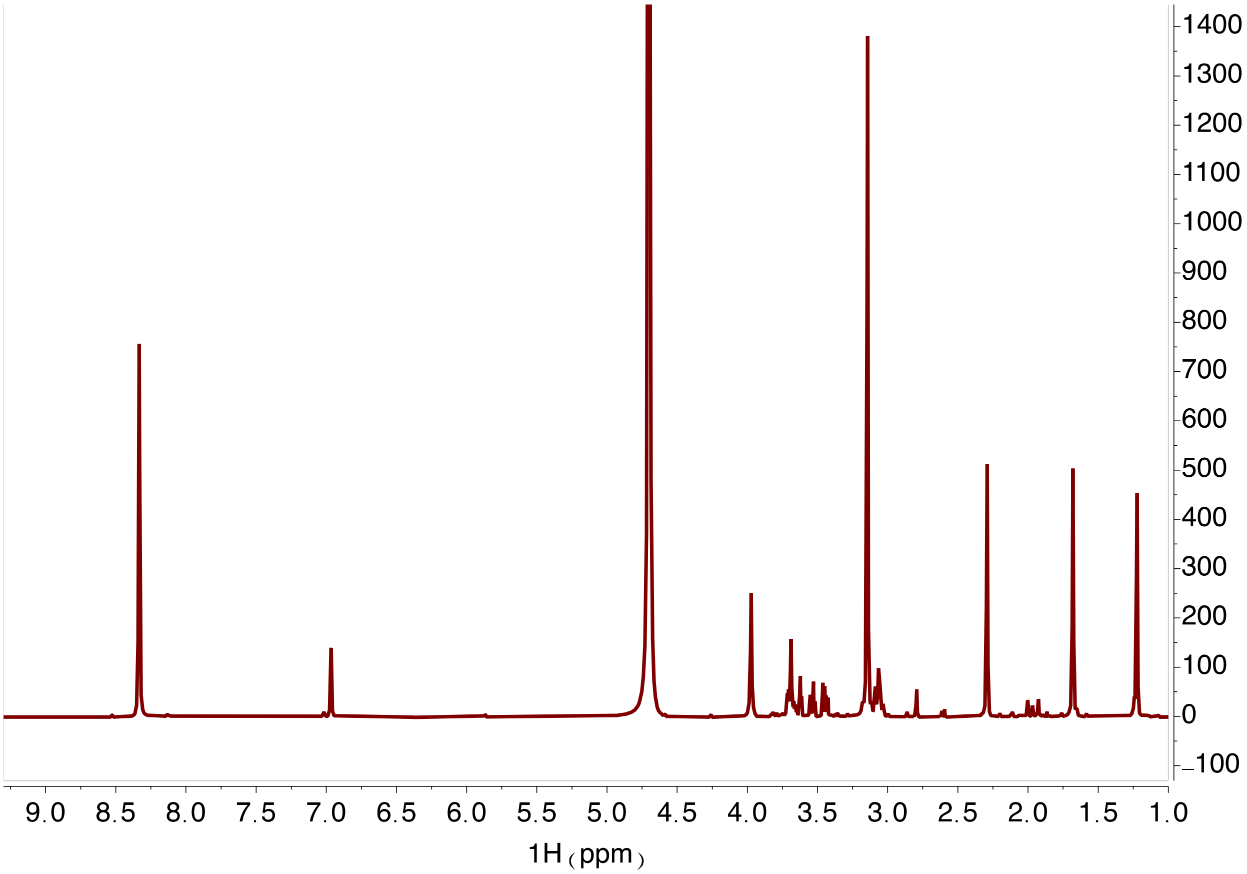

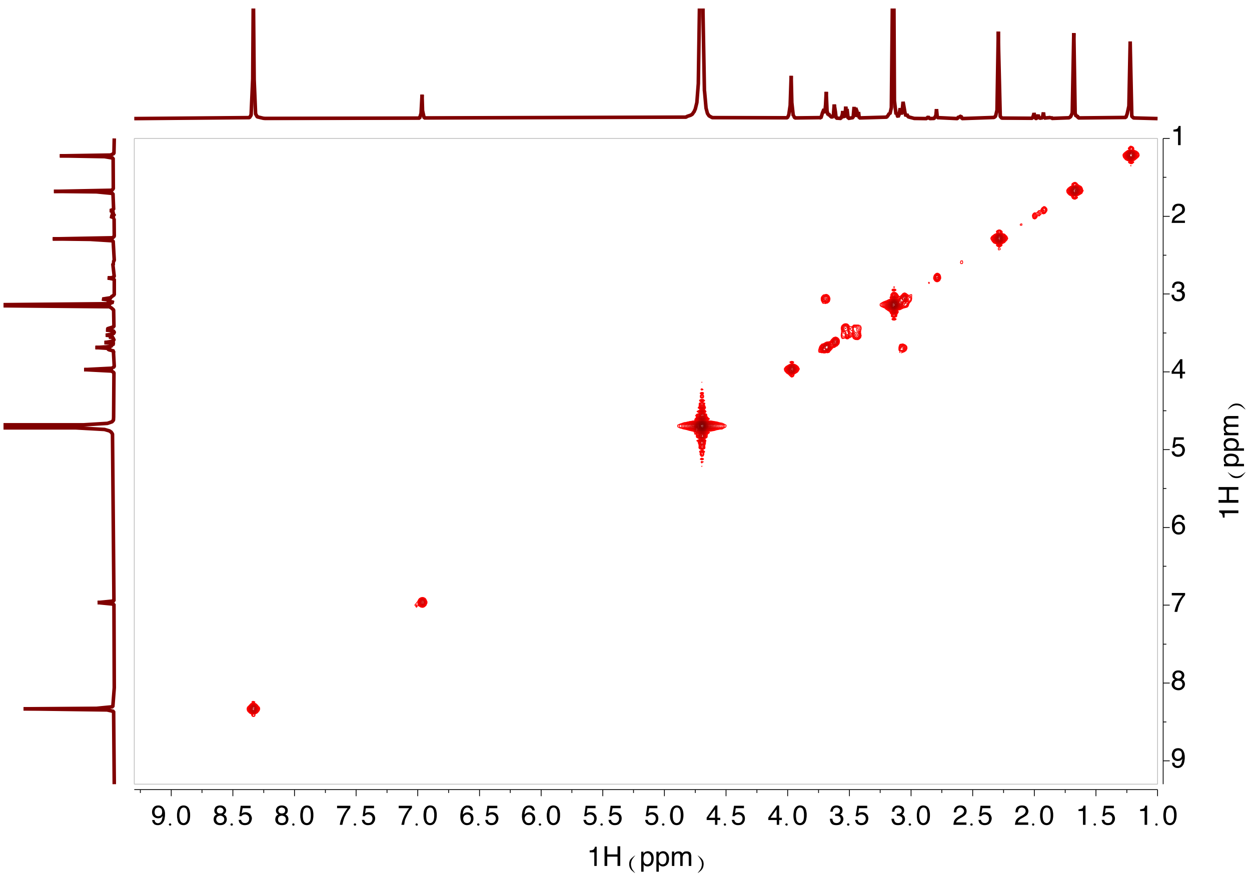
9.0 8.5 8.0 7.5 7.0 6.5 6.0 5.5 5.0 4.5 4.0 3.5 3.0 2.5 2.0 1.5 1.0
1H（ppm）
1 2 3 4 5 6 7 8 9
1H（ppm）

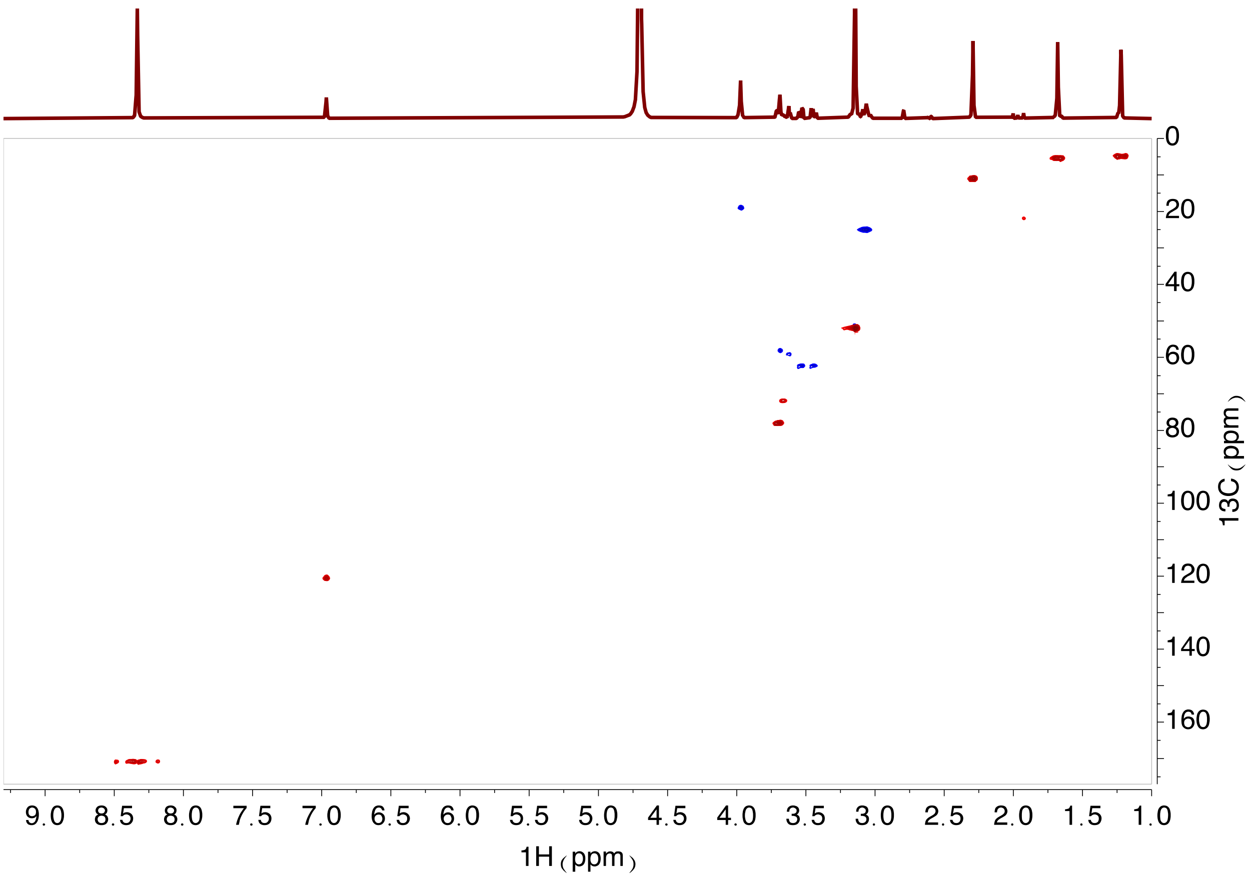
9.0 8.5 8.0 7.5 7.0 6.5 6.0 5.5 5.0 4.5 4.0 3.5 3.0 2.5 2.0 1.5 1.0
1H（ppm）
0 20 40 60 80 100 120 140 160
13C（ppm）

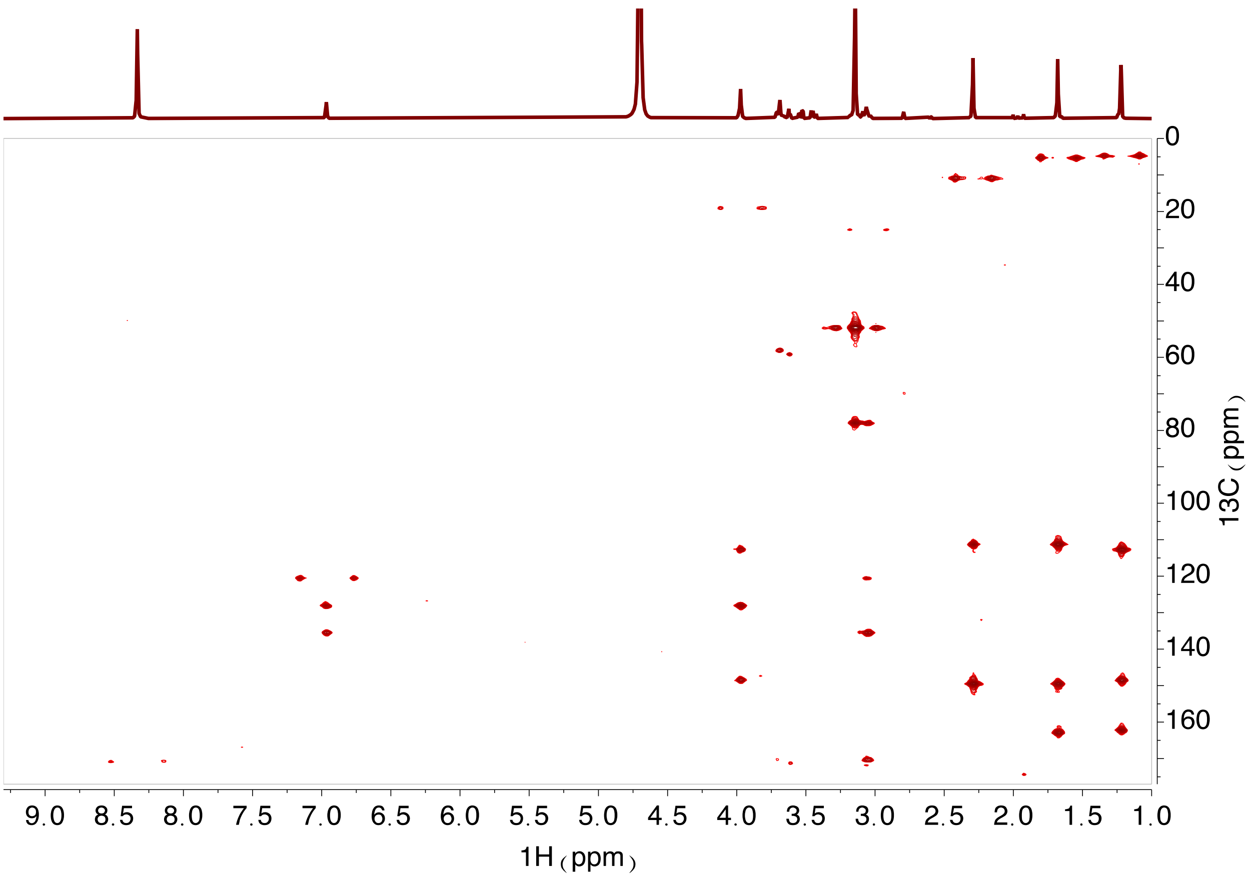
9.0 8.5 8.0 7.5 7.0 6.5 6.0 5.5 5.0 4.5 4.0 3.5 3.0 2.5 2.0 1.5 1.0
1H (ppm)
0 20 40 60 80 100 120 140 160
13C (ppm)

**Supplementary Figure 9.** HR-MS/MS profiles for products of the SenA reaction. The structure of each product (i.e., parent ion) is shown to the right. The predicted identities of the numbered fragments are displayed in Supplementary Table 8.

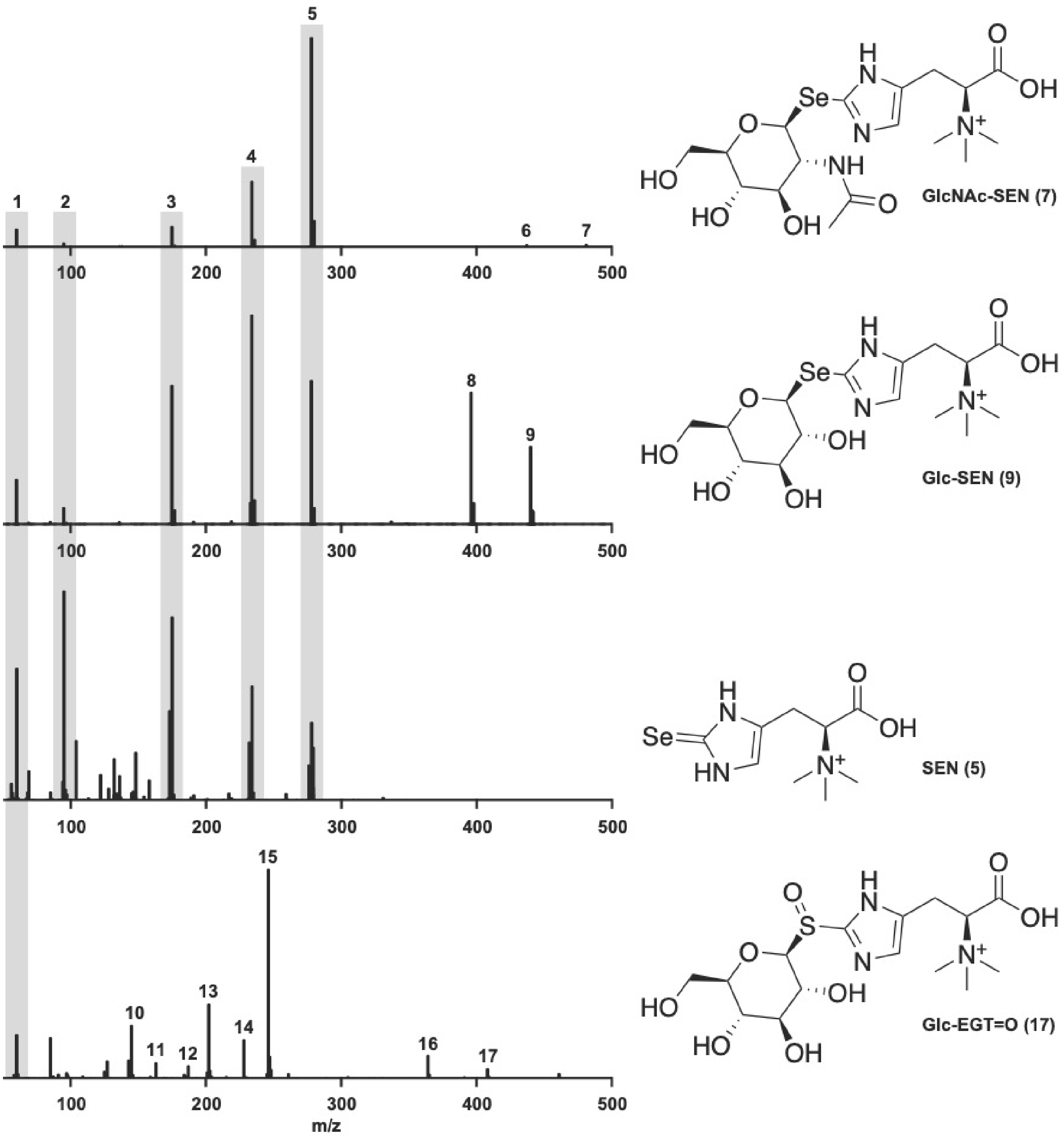
